# Early-stage sparse testing strategies to increase genetic gain in plant breeding programmes

**DOI:** 10.1101/2025.08.18.670883

**Authors:** Christian R Werner, Giovanny Covarrubias-Pazaran, David González-Diéguez, Dorcus C Gemenet, Harish Gandhi, Gary N Atlin

## Abstract

Early-stage sparse testing can significantly increase genetic gain in plant breeding programmes, facilitating the development of varieties with high and stable performance across farmers’ fields. This is achieved through (1) increased selection accuracy, enabled by broader sampling of trial sites across the Target Population of Environments (TPE), and (2) increased selection intensity by testing more selection candidates at a reduced replication rate.

Early-stage agronomic testing typically involves evaluating a large number of selection candidates at one or a few experimental sites. Although these sites may poorly represent the TPE, strong selection pressure is generally applied. However, if the genetic correlation between performance at the experimental sites and in the TPE is low, most of the genetic gain achieved through selection will not be expressed under farmers’ conditions. Sparse testing addresses this challenge by using farm-as-incomplete-block designs, in which different subsets of selection candidates are evaluated across sites. By leveraging a genomic relationship matrix (GRM) to connect genotypes across environments, sparse testing enables early-stage multi-environment trials with broader coverage of the TPE.

Here, we used stochastic simulation to compare various partially replicated sparse testing strategies to three fully replicated conventional early-stage testing strategies, with and without GRM. All sparse testing strategies achieved substantially higher and more stable genetic gain than the conventional strategies. Our results show that sparse testing provides breeders with a powerful and flexible framework to rethink early-stage trials and design cost-effective, multi-location experiments that lay the foundation for increased genetic gains in farmers’ fields.

**Key message:** Early-stage sparse testing can significantly increase genetic gain by (1) improving selection accuracy through broader sampling of the Target Population of Environments (TPE), and (2) increasing selection intensity.

## Introduction

Plant breeding programmes aim to develop varieties that exhibit high and stable performance across a systematically delineated range of growing conditions, often spanning diverse production environments. The entire spectrum of crop production environments targeted by a breeding programme is referred to as the target population of environments (TPE). The TPE includes all farms and future seasons in which the varieties produced by a breeding programme will be grown (Comstock, 1977; Chenu et al., 2011; Werner et al., 2025) and, therefore, is the environment in which genetic gain is desired (van Eeuwijk et al., 2001). It is characterized by its biophysical conditions, encompassing both biotic and abiotic environmental factors, as well as the agronomic management practices adopted by farmers (Ceccarelli, 1989, 1994; Chenu et al., 2011; Cooper et al., 2020, 2021, 2022). As such, the TPE is a critical concept in plant breeding which recognizes the variation in environments where crops are grown (Cooper and DeLacy, 1994; Cooper et al., 1996, 2021; van Eeuwijk et al., 2001). It allows breeding organisations and seed companies to align their logistical and commercial constraints with the demands and preferences of farmers, processors, and end-users (Atlin et al., 2000), captured within the concept of a market segment (Covarrubias-Pazaran et al., 2022).

To identify the best parents for population improvement and candidate varieties for product development (Gaynor et al., 2017), breeders must test germplasm targeting a specific market segment under conditions that closely resemble those encountered in farmers’ fields (van Eeuwijk et al., 2001; Ceccarelli and Grando, 2009; Ceccarelli, 2015; Cooper et al., 2016). However, accomplishing this is particularly challenging during the early stages of agronomic testing in breeding trials, which typically involve evaluating a large number of selection candidates with limited multiplication capacity (Ceccarelli, 2015), thus restricting the programme’s ability to conduct multi-environment trials that capture the full range of conditions within the TPE. Given the logistical challenges of testing hundreds or thousands of selection candidates and ensuring comparability among them, early-stage trials are usually conducted at only a few experimental sites managed by breeders. However, these experimental sites are, by definition, not part of the TPE. To address this, breeders typically aim to select sites within the same geographical regions, artificially reproduce conditions such as drought or heat stress, and incorporate common management practices in their trials (Werner et al., 2025). Despite these efforts, limited resources often lead breeders to concentrate on only a subset of environmental factors, meaning their trials may not fully reflect the wide range of conditions farmers will face. Moreover, while phenotyping on experimental sites typically focuses on individual stress factors in isolation, farmers’ fields often experience combinations of multiple biotic and abiotic stresses (Cooper and Messina, 2021, 2023). Resilience to these combined stresses is complex and may not be directly predictable from a genotype’s response to each stress factor in isolation, and determining how to integrate and prioritize individual measurements into a predictor of genetic gain in the TPE often remains unclear. While this is generally a challenge in plant breeding, it is particularly pronounced in public breeding programmes due to the diverse and extensive range of environments they often aim to serve.

Despite those limitations during early-stage trials in many breeding pipelines, substantial selection pressure is applied particularly in the initial agronomic trials. In most CGIAR breeding programmes, for example, well over 90% of the selection pressure is concentrated at a few managed research stations, where inferior candidates are eliminated in unreplicated nurseries or initial low-replication agronomic trials, typically based on visual selection (Ceccarelli, 1989, 2015; Ceccarelli and Grando, 2009). This strategy assumes that rapidly and cost-effectively culling large populations will allow for a more efficient allocation of testing resources to a superior subset of candidates in later stages.

However, the combination of high selection intensity with testing at only a few sites during early-stage trials carries the risk that these sites may not accurately reflect the TPE, potentially enriching haplotypes in the population that may not translate into improved performance on farmers’ fields. If the genetic correlation between performance on managed experimental sites and in the TPE is low, only a small portion of the genetic gain measured at those sites will be carried on to later stages of testing and expressed in farmers’ fields (Atlin et al., 2001; Cooper et al., 2023). This highlights the need for a strategy to evaluate early-stage material across a wide range of locations that capture the diversity of conditions in the TPE.

Sparse testing, a multi-environment strategy in which not all selection candidates are grown in every testing environment (Endelman et al., 2014), presents an opportunity to fundamentally rethink early-stage testing designs in plant breeding programmes. It employs farm-as-incomplete-block (FAIB) designs (Atlin et al., 2001; Werner et al., 2025), where only a subset of selection candidates, and therefore of haplotype combinations, is tested at each site. This approach is similar to genetic evaluation methods used in smallholder dairy farms (Powell et al., 2021). The implementation of sparse testing is enabled by the use of a genomic relationship matrix (GRM), which leverages genomic data to connect unreplicated selection candidates, and enable fair and meaningful comparisons across multiple trial sites (Endelman et al., 2014; Werner et al., 2025). Information on unreplicated selection candidates is enhanced through observations on numerous closely related family members (Windhausen et al., 2012; Lorenz, 2013), effectively serving as ’partial replicates’. This approach allows for multi-environment trials with an expanded representation of the TPE, providing a more robust evaluation of the breeding population under real-world conditions at the earliest stage of agronomic testing when selection pressure is highest.

We hypothesize that sparse testing strategies have significant potential to optimize early-stage testing in plant breeding programmes and increase genetic gain in farmers’ fields. First, by expanding the number of test locations, sparse testing allows the population to be evaluated across a much broader range of conditions, thereby increasing selection accuracy within the TPE. Second, by decreasing individual replication rates, sparse testing optimizes resource use and enables breeders to evaluate more individuals and, thus, a greater diversity of haplotype combinations. This not only expands the size of the training population for genomic prediction models but also increases selection intensity, thereby strategically addressing two components of the breeder’s equation (Lush, 1937) simultaneously.

To test our hypothesis, we used stochastic simulation to compare seven partially replicated sparse testing strategies to three fully replicated conventional early-stage testing strategies with and without incorporation of a GRM in the trial analysis. We found that sparse testing resulted in substantially higher and more stable rates of genetic gain compared to conventional strategies. Thus, sparse testing provides breeders with greater flexibility to design cost-effective, multi-location early-stage trials, laying the foundation for extending early-stage testing to farmers’ fields.

## Methods

Stochastic simulation was used to evaluate the potential of sparse testing designs to optimize early-stage testing in plant breeding programmes. Therefore, seven partially replicated sparse testing strategies were compared to three fully replicated conventional early-stage testing strategies with and without incorporation of a genomic relationship matrix (GRM) in the trial analysis. We simulated a quantitative trait, such as yield, and assessed the testing strategies for genetic gain in the target population of environments (TPE), stability of genetic gain, and prediction accuracy over 10 cycles of crossing and selection.

The methods are subdivided into three sections. The first section describes the simulation of the founder population. The second section describes the simulation of the early-stage testing strategies. The third section describes the evaluation and comparison of the testing strategies.

### Simulation of the founder population

#### Genome simulation

Whole-genome sequences were simulated for a founder population of 30 homozygous genotypes of a diploid line crop species. Each founder genome consisted of 10 chromosome pairs with a physical length of 2 x 10^8^ base pairs (bp) and a genetic length of 200 centimorgans (cM), resulting in a total physical length of 2 Gbp and a genetic length of 2 Morgan. Whole-genome sequences were generated using the Markovian coalescent simulator (Chen et al., 2009) in AlphaSimR (Gaynor et al., 2021), deploying AlphaSimR’s maize evolutionary history default settings. Recombination rate was derived as the ratio between genetic length and physical length (i.e., 2 Morgan / 2x10^8^ bp = 10^-8^ recombinations per base pair), and mutation rate was set to 1.25 x 10^-8^ per base pair. The effective population size (N_e_) at the end of the coalescent process was set to 100. A set of 600 biallelic quantitative trait nucleotides (QTN) and 300 single nucleotide polymorphisms (SNP) was randomly sampled along each chromosome to simulate a quantitative trait controlled by 6,000 QTN and a SNP marker array with 3,000 markers. Note that the coalescence process was primarily intended to generate linkage disequilibrium (LD) in the founders (Werner et al., 2023), rather than to create a population representative of maize as a species. Beyond LD generation, the coalescence step generally has negligible effects on overall simulation results.

#### Simulation of the Target Population of Environments (TPE)

A total of 2,000 locations were simulated by creating correlated genetic values across locations for the 30 genotypes in the founder population, using an unstructured model for genotype-by-environment (GxE) interaction (van Eeuwijk et al., 2001; Smith et al., 2005; Malosetti et al., 2013). These locations represent the TPE defined and targeted by a breeding programme, such as a geographical region assumed to share similar biophysical conditions and agronomic management practices. However, due to permanent factors, such as suboptimal management, or temporary factors, like extreme weather conditions, data collected at some sites may not be representative of the TPE. To account for this, the simulated locations were randomly divided into two sets: a larger set of locations representative of the TPE and a smaller set, not representative of the TPE, referred to as off-TPE locations. The number of off-TPE locations was resampled for each of 100 simulation replicates, ranging from 100 to 200 (i.e., 5-10% of the total number of locations were not representative of the TPE).

A 2000 x 2000 matrix of pairwise genetic correlations between locations was generated for each simulation replicate using the ‘*struc_cor_mat*’ function in FieldSimR (Werner et al., 2024), with arguments specified as follows: baseline genetic correlations were set to 0.6 among TPE locations, 0.4 among off-TPE locations, and -0.5 between TPE and off-TPE locations; the range of correlations around the baseline values was set to 0.3; and the rank of the correlation matrix was set to 13. These settings produced distributions of pairwise genetic correlations centred around a mean of approximately 0.6 within the TPE and off- TPE groups, and approximately -0.5 between TPE and off-TPE locations. To illustrate this, Figure 1 shows the distribution of pairwise genetic correlations between locations for one of the simulation replicates. This correlation structure resulted, on average, in a ratio of 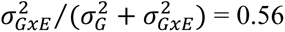, with 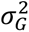 representing the genetic main effect variance across all simulated environments and 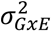 the variance due to interaction not explained by the main effect. Across the 100 simulation replicates, crossover interaction variance contributed, on average, 53% of the total genetic variance, while non-crossover variance contributed 47% (Tab. S2).

**Figure 1.**
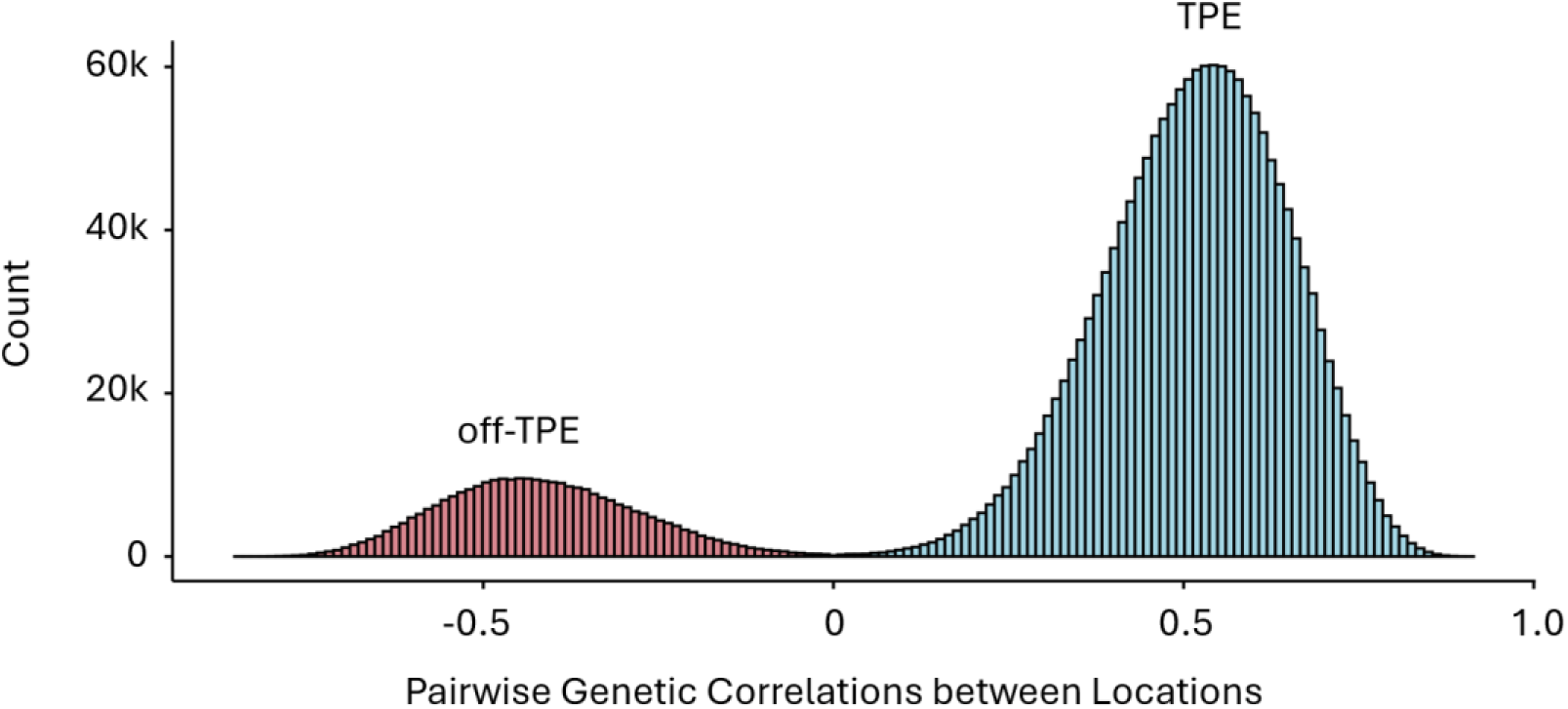
Histogram of pairwise genetic correlations between the 2,000 simulated locations representing the target population of environments defined and targeted by a breeding programme in one of the simulation runs. The blue distribution (TPE) depicts the set of locations that are representative of the target population of environments. The red distribution (off-TPE) depicts the set of locations that are not representative of the target population of environments.

Note that we chose these values to reflect what we consider, based on our experience, to be a reasonably well-defined TPE suitable for a single breeding pipeline. We acknowledge that some real-world datasets will exhibit lower pairwise genetic correlations among locations and/or include a higher proportion of off-TPE sites. However, substantially lower correlations or a consistently large number of off-TPE locations may indicate that the TPE is too diverse to be addressed by a single pipeline, limiting the breeder’s ability to identify genotypes with consistently strong performance across all environments. Such a scenario would call for redefining the TPE, rather than simply improving the sampling strategy.

Location-specific additive genetic effects (*a*) were sampled from a multivariate normal distribution, employing the 2000 x 2000 matrix of pairwise genetic correlations. Genetic values for a single quantitative trait, such as yield, were simulated by summing the additive genetic effects at the 6,000 QTN and scaled to obtain a location-specific additive genetic variance 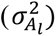. Mean genetic values of the population were sampled for each location individually from a standard normal distribution. Genetic variances were first sampled from a chi-squared distribution with degrees of freedom set to 100, and then divided by 100 to obtain a mean additive genetic variance in the founder population of approximately 1 across all 2,000 locations, i.e., 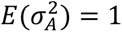. The true genetic values of the 30 founder genotypes were then calculated as the mean of the genetic values across all locations representative of the TPE (i.e., off-TPE locations were excluded).

#### Simulation of phenotypes

Phenotypes were generated by adding random error to the location-specific genetic values. The random error was sampled from a normal distribution with a mean of zero and a location-specific error variance 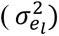 derived from the equation for narrow-sense heritability, assuming a plot-level heritability of ℎ^2^ = 0.1 in the founder population across all locations:

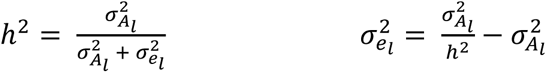

With the location-specific error variance held constant throughout crossing cycles, the decrease in genetic variance due to selection (Fig. S5) also led to a gradual decline in heritability (ℎ^2^) over time.

### Simulation of the early-stage testing strategies

#### Simulation scenarios

We simulated 10 cycles of crossing, testing, and parent selection in a diploid line crop for each of seven partially replicated sparse testing strategies, and three fully replicated conventional early-stage testing strategies with and without incorporation of a GRM in the trial analysis. This resulted in a total of 13 simulation scenarios, presented in Table 1. At each cycle, 30 inbred parents were selected and randomly crossed to create the next generation of inbred lines. While early-stage testing resources were fixed at 2,200 plots in all simulation scenarios, we systematically varied the number of crosses, genotypes per cross, total genotypes, test sites, plots per site, and replication rates across scenarios. Our objective was to explore how testing strategies can be optimised under fixed plot constraints. At each cycle, the test sites were randomly sampled from the 2000 simulated locations. All simulation scenarios were replicated 100 times.

**Table 1.** Simulated early-stage testing scenarios.

| Conventional Testing Scenarios |  |  |  |  |  |  |  |
| --- | --- | --- | --- | --- | --- | --- | --- |
| Scenario* | Crosses | Lines / cross | Lines tested | Test sites | Reps / site | Plots / site | GRM |
| CT-550_2 | 22 | 25 | 550 | 2 | 2 | 1100 | No |
| CT-440_5 | 22 | 20 | 440 | 5 | 1 | 440 | No |
| CT-220_5 | 22 | 10 | 220 | 5 | 2 | 440 | No |
| CT-550_2-GRM | 22 | 25 | 550 | 2 | 2 | 1100 | Yes |
| CT-440_5-GRM | 22 | 20 | 440 | 5 | 1 | 440 | Yes |
| CT-220_5-GRM | 22 | 10 | 220 | 5 | 2 | 440 | Yes |

| Sparse Testing Scenarios |  |  |  |  |  |  |  |
| --- | --- | --- | --- | --- | --- | --- | --- |
| Scenario* | Crosses | Lines / cross | Lines tested | Test sites | Reps (in TPE) | Plots / site | GRM |
| ST-5_440 | 40 | 50 | 2000 | 5 | 0.1 | 440 | Yes |
| ST-10_220 | 40 | 50 | 2000 | 10 | 0.1 | 220 | Yes |
| ST-20_110 | 40 | 50 | 2000 | 20 | 0.1 | 110 | Yes |
| ST-50_44 | 40 | 50 | 2000 | 50 | 0.1 | 44 | Yes |
| ST-100_22 | 40 | 50 | 2000 | 100 | 0.1 | 22 | Yes |
| ST-200_11 | 40 | 50 | 2000 | 200 | 0.1 | 11 | Yes |
| ST-440_5 | 40 | 50 | 2000 | 440 | 0.1 | 5 | Yes |
| BSLN-1_2200 | 40 | 50 | 2000 | 1 | 0.1 | 2200 | Yes |

**\*Note:** The numbers in the names of the conventional testing (CT) scenarios represent the total number of genotypes tested and test sites, while the numbers in the names of the baseline (BSLN) and the sparse testing (ST) scenarios correspond to the number of test sites and plots per site.

Each simulation scenario was preceded by a burn-in period consisting of three cycles of random crossing and selection, executed as in the conventional testing scenario CT-550_2 (Tab. 1). This burn-in period was used to establish a base level of family structure in the breeding population, which is essential for genomic prediction to be effective in breeding populations. The final burn-in cycle served as a common starting point for all simulation scenarios.

In the first burn-in cycle, the 30 homozygous founder genotypes served as parents. In the following two cycles, 30 new inbred parents were randomly selected from the current generation, ensuring that each parent came from a different cross. These parents were then randomly combined to generate 22 unique bi-parental crosses. Each cross produced 25 heterozygous F_1_-derived progeny genotypes, resulting in a population of 550 genotypes. From each progeny genotype, one fully homozygous line was derived to mimic a single-seed descent breeding scheme. Parents were selected at random to keep the genetic variance somewhat stable during the burn-in period. The parents selected in the last burn-in cycle served as initial parents for the 13 testing scenarios.

#### Simulation of the conventional early-stage testing strategies

The three conventional early-stage testing strategies (CT-550_2, CT-440_5, CT- 220_5) represent typical stage 1 and stage 2 testing strategies conducted by breeders on managed experimental stations, similar to those currently employed by most CGIAR breeding programmes. Since the entire set of genotypes is tested at least once at each location (Tab. 1), the conventional early-stage testing strategies are characterised by a trade-off between the number of genotypes, the number of test sites, and the number of replicates per site. Each strategy provides a different interpretation of this trade-off. The three strategies were simulated both with and without a GRM included in the trial analysis to connect genotypes across sites.

#### Simulation of the sparse-testing strategies

The seven sparse testing (ST) strategies implemented a farm-as-incomplete-block (FAIB) design (Atlin et al., 2001; Werner et al., 2025) to enable early-stage testing across a larger number of test sites while keeping the total number of plots constant. Combined with a partial replication rate of 0.1 across all test sites, the FAIB sparse testing designs allowed for a substantial increase in the total number of genotypes compared to the conventional early-stage testing strategies. Specifically, of the 2,000 tested genotypes, 1,800 were unreplicated, while 200 were replicated once. Note that the partial replication rate reflects replication across all test sites, with replication within individual sites deliberately avoided.

All sparse testing strategies used the same number of crosses, genotypes per cross, and total number of genotypes. The genotypes were distributed across an increasing number of sites (Tab. 1). Allocation to test sites was performed using a restricted randomization algorithm, which minimized the occurrence of full-sibs at the same site to ensure an even distribution of families across locations (available on: https://github.com/crWerner/sparseTesting). With a fixed total number of 2,000 genotypes and a partial replication rate of 0.1, the sparse testing strategies must optimize the trade-off between the number of test sites and the number of plots per site.

A baseline scenario (BSLN-1_2200) was also included, in which all 2,000 genotypes and their partial replicates were tested at a single site, representing an early-generation nursery grown at only one location. The baseline scenario served as a reference point for evaluating the effect of distributing selection candidates across multiple sites. All sparse testing strategies utilized a GRM, which is essential for connecting unreplicated genotypes across incomplete blocks (sites) and ensuring comparability between the tested genotypes.

### Comparison of the early-stage testing strategies

To compare the sparse testing and conventional early-stage testing strategies, we assessed genetic gain, stability of genetic gain, and selection accuracy across 10 cycles of crossing, testing, and selection. Genetic gain was measured as the mean genetic value of the population across all locations that were representative of the TPE. Stability of genetic gain was evaluated based on the standard deviation and coefficient of variation of genetic gain across the 100 simulation replicates. Selection accuracy was determined by the Pearson correlation coefficient between the true genetic values of the genotypes in the TPE and the breeding values or genomic estimated breeding values (GEBV), depending on whether the GRM was included in the analysis.

#### Analysis of the early-stage testing data

The prediction of (genomic estimated) breeding values for selection of parents was done using the following linear mixed model:

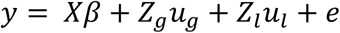

Where 𝑦 is the vector of phenotypic observations, 𝛽 is a vector of the fixed overall mean effect with design matrix 𝑋, 𝑢_𝑔_ is a vector of random (genomic estimated) breeding values with design matrix 𝑍_𝑔_, 𝑢_𝑙_ is a vector of random location effects with design matrix 𝑍_𝑙_, and 𝑒 is the vector of residuals. We assumed random effects to be normally distributed:

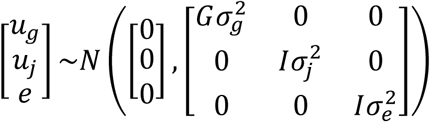

With 𝐺 being the genomic relationship matrix as proposed by vanRaden (2008) when included in the analysis, or 𝐺 = 𝐼 when no genomic relationship matrix was included. Linear mixed model analyses to calculate best linear unbiased predictors (BLUP) of the breeding values were performed using the sommer R package version 4.1.0 (Covarrubias-Pazaran, 2016).

## Results

Our results demonstrate that early-stage sparse testing strategies can significantly increase genetic gain in plant breeding programmes compared to conventional fully replicated testing strategies. This improvement is achieved through an efficient combination of high selection accuracy, facilitated by broader sampling of locations for testing across the target population of environments (TPE), and increased selection intensity.

In comparison to the conventional early-stage testing strategies which incorporated a genomic relationship matrix (GRM) in the trial analysis, sparse testing could increase genetic gain by more than 20% after 10 cycles of crossing, testing, and selection. In contrast, the inclusion of a GRM in the conventional testing strategies led to a 7% to 24% increase in genetic gain compared to when no GRM was used. Since all seven sparse testing strategies produced relatively similar results, we present only three selected simulation scenarios: the strategy with the highest (ST-50) and lowest (ST-5_440) average genetic gain, as well as the strategy with the most sites (ST-440_5). Summary statistics comparing genetic gain and gain stability across the simulated conventional early-stage testing scenarios, the three sparse testing scenarios, and the baseline scenario are presented in Table 2. The results for all other sparse testing strategies are provided in the supplementary material.

**Table 2.** Summary statistics for the simulated conventional early-stage testing scenarios, the three selected sparse testing scenarios, and the baseline scenario. The table includes average genetic gain (Gain), minimum (Min) and maximum (Max) genetic gain, standard deviation of genetic gain (SD), and coefficient of variation of genetic gain (CV), based on 100 simulation replicates.

| Scenario | Gain | Min | Max | SD | CV |
| --- | --- | --- | --- | --- | --- |
| CT-220_5 | 3.28 | 2.04 | 5.63 | 0.69 | 20.97 |
| CT-440_5 | 3.06 | 1.79 | 5.71 | 0.74 | 24.05 |
| CT-550_2 | 2.84 | 1.55 | 4.40 | 0.64 | 22.60 |
| CT-220_5-GRM | 3.51 | 1.92 | 5.21 | 0.64 | 18.21 |
| CT-440_5-GRM | 3.67 | 2.61 | 6.08 | 0.66 | 18.02 |
| CT-550_2-GRM | 3.51 | 1.31 | 5.28 | 0.73 | 20.72 |
| ST-5_440 | 4.32 | 2.71 | 6.80 | 0.74 | 17.11 |
| ST-50_44 | 4.48 | 3.20 | 6.42 | 0.71 | 15.79 |
| ST-440_5 | 4.37 | 3.03 | 6.05 | 0.65 | 14.84 |
| BSLN-1_2200 | 3.83 | 1.11 | 6.4 | 0.91 | 23.88 |

### Genetic gain in the target population of environments (TPE)

All sparse testing strategies generated substantially greater genetic gain than the conventional early-stage testing strategies. This is shown in Figure 2, which plots genetic gain in the TPE against time for the three sparse testing (ST) strategies (solid lines), the three conventional early-stage testing (CT) strategies with (solid lines) and without GRM (dashed lines), and the baseline scenario (dotted lines). For example, the best sparse testing strategy (ST-50_44) generated 22% more genetic gain than the best conventional strategy with GRM (CT-440_5-GRM), and 37% more gain than the best conventional strategy without GRM (CT-220_5).

**Figure 2.**
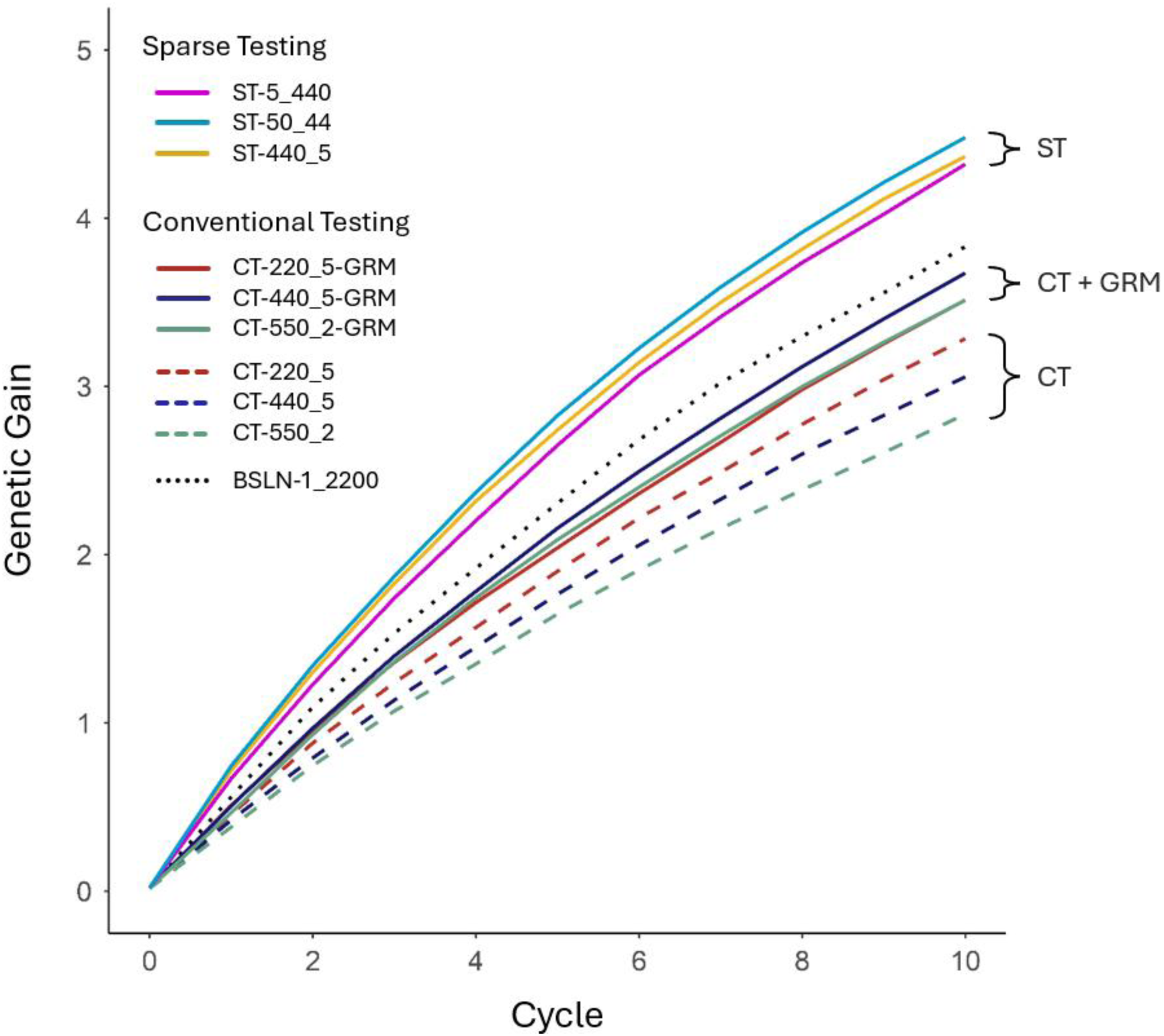
Genetic gain of the simulated conventional early-stage testing strategies without (CT-220_5, CT-440_5, CT-550_2) and with a genomic relationship matrix (CT-220_5-GRM, CT-440_5-GRM, CT-550_2-GRM), the sparse testing strategies (ST-5_440, ST-50_44, ST-440_5), and the baseline scenario (BSLN-1_2200) which represent a single-site early generation nursery including a GRM. Genetic gain is plotted as the mean genetic value of the population across all locations representative of the target population of environments. Each line shows the change in mean genetic value over 10 cycles of crossing, testing, and selection for 100 simulated replicates.

Figure 2 also shows that the difference in genetic gain between the best (ST-50_44) and worst (ST-5_440) sparse testing strategies was marginal. After 10 cycles of crossing, testing, and selection, the ST-50_44 strategy achieved only 4% more genetic gain than the ST-5_440 strategy, highlighting that all sparse testing strategies performed comparably in terms of average genetic gain (Tab. S1, Fig. S1). In contrast, the baseline strategy (BSLN- 1_2200), conducted at only one site, yielded 15% less genetic gain than the ST-50_44 strategy. Nevertheless, on average, it still generated 4% more gain than the best conventional testing strategy with GRM, due to the moderate accuracy and higher selection intensity it afforded.

Figure 2 also shows that the inclusion of a GRM in the analysis of conventional early-stage trials consistently enhanced genetic gain across all three scenarios. Specifically, in cycle 10, this resulted in a 7% increase in genetic gain in CT-220_5-GRM, 20% in CT- 440_5-GRM, and 24% in CT-550_2-GRM. Furthermore, including a GRM led to a reranking of the three conventional testing strategies. Without the GRM, the CT-220_5 strategy achieved the highest genetic gain, generating 7% more gain than CT-440_5 and 15% more than CT-550_2. However, with the GRM included, the CT-440_5-GRM strategy emerged as the most effective, achieving approximately 4% more gain than both CT-220_5-GRM and CT-550_2-GRM, which showed similar gains after 10 cycles.

### Stability of genetic gain

Early-stage sparse testing designs with low replication offer high rates of genetic gain coupled with increased gain stability. This is shown in Figure 3, which plots genetic gain against time for all 100 simulation replicates for two of the conventional early-stage testing strategies (CT-220_5 and CT-550_2), both with and without the GRM (top), as well as the three sparse testing strategies and the baseline strategy (bottom). Among the sparse testing strategies, testing in more sites resulted in greater stability of genetic gain across the 100 simulation replicates.

**Figure 3.**
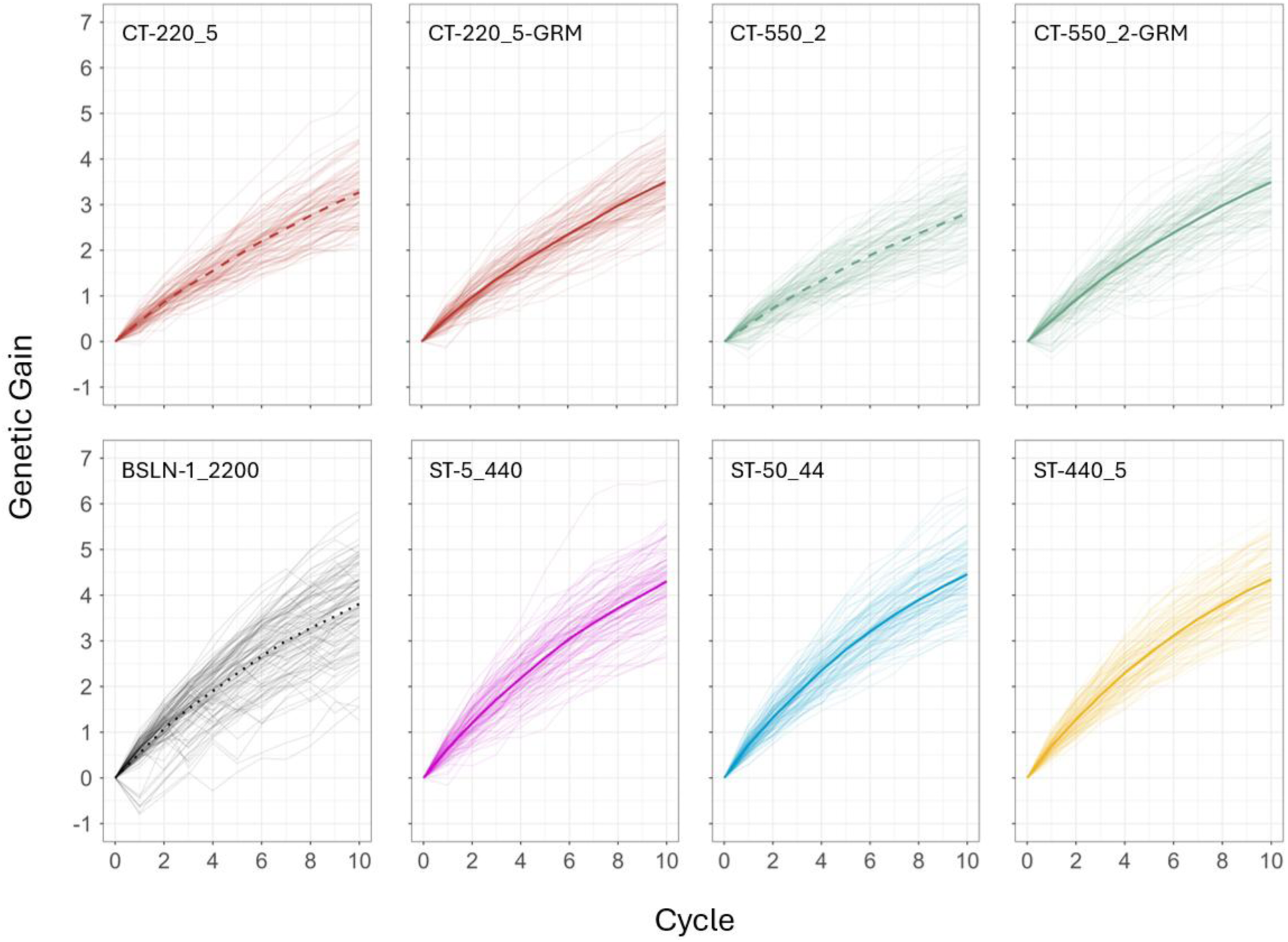
Genetic gain of each of the 100 simulation replicates of two of the conventional early-stage testing strategies without (CT-220_5, CT-550_2) and with a genomic relationship matrix (CT-220_5-GRM, CT-550_2-GRM), the sparse testing strategies (ST-5_440, ST-50_44, ST-440_5), and the baseline scenario (BSLN-1_2200). Genetic gain is shown as the mean genetic value of the population across all locations representative of the target population of environments. Each line represents the change in genetic value over 10 cycles of crossing, testing, and selection, with bold lines showing the mean across the 100 replicates.

Specifically, ST-50_44 and ST-440_5 not only achieved on average slightly more genetic gain than ST-5_440 (Fig. 2 & 3) but also showed lower standard deviation (Fig. 4a) and a lower coefficient of variation (Fig. 4b) of genetic gain over time. Moreover, the coefficients of variation in gain over time of the sparse testing scenarios were lower than those of the conventional testing scenarios, both with and without GRM.

**Figure 4.**
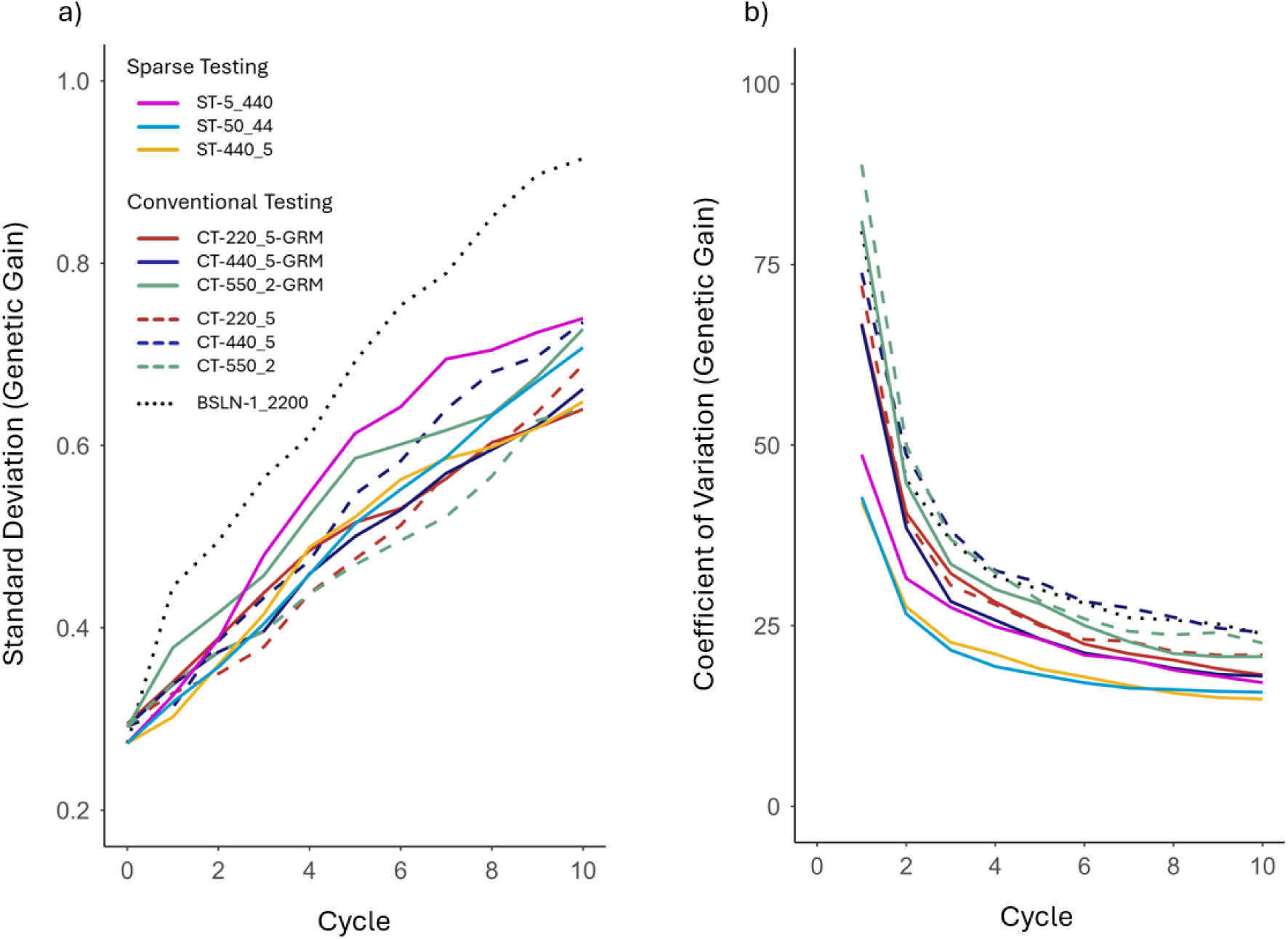
Standard deviation and coefficient of variation of genetic gain for the simulated conventional early-stage testing strategies without (CT-220_5, CT-440_5, CT-550_2) and with a genomic relationship matrix (CT-220_5-GRM, CT-440_5-GRM, CT-550_2-GRM), the sparse testing strategies (ST-5_440, ST-50_44, ST-440_5), and the baseline scenario (BSLN-1_2200). Each line shows the change in standard deviation (a) and coefficient of variation (b) over 10 cycles of crossing, testing, and selection. To aid visualization, coefficient of variation values at cycle 0 were omitted, as they ranged between 1,300 and 1,800. The standard deviation and coefficient of variation were calculated from genetic gain across the 100 simulation replicates.

The highest coefficient of variation, however, was observed in the CT-440_5 scenario, which achieved slightly more genetic gain than the CT-550_2 strategy but also showed a higher standard deviation. As with genetic gain, the CT-440_5 strategy benefitted the most from the inclusion of a GRM (CT-440_5-GRM) among the conventional testing strategies, as demonstrated by a substantial reduction in both the standard deviation (Fig. 4a) and the coefficient of variation (Fig. 4b) in gain.

### Selection accuracy

Sparse testing selection accuracies were in a range comparable to the selection accuracies of the conventional early-stage testing strategies that included a GRM. This is shown in Figure 5, which plots selection accuracy of the (genomic estimated) breeding values in the TPE against time for the three conventional early-stage testing strategies (Fig. 5a) with (solid lines) and without a GRM (dashed lines), as well as the three sparse testing strategies (solid lines) and the baseline scenario (dotted lines) (Fig. 5b). To aid visualization and the comparison between the conventional and the sparse testing strategies, Figure 5 is divided into two panels.

**Figure 5.**
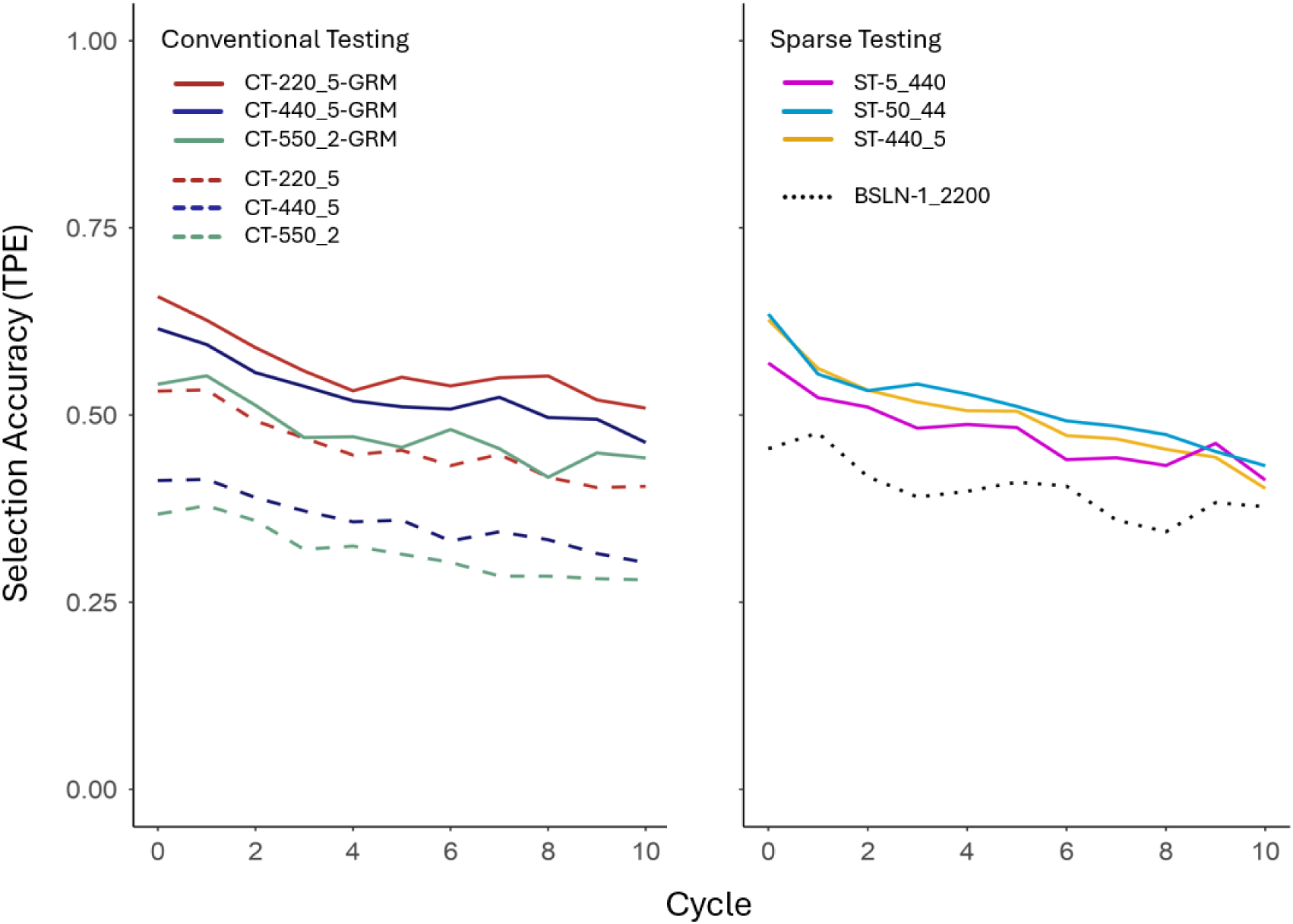
Selection accuracy for the simulated conventional early-stage testing strategies without (CT-220_5, CT-440_5, CT-550_2) and with a genomic relationship matrix (CT-220_5-GRM, CT-440_5-GRM, CT-550_2-GRM), the sparse testing strategies (ST-5_440, ST-50_44, ST-440_5), and the baseline scenario (BSLN-1_2200). Selection accuracy was calculated as the Pearson correlation coefficient between the genetic values of the genotypes across all locations representative of the target population of environments and their breeding values or genomic estimated breeding values (GEBV), depending on whether the genomic relationship matrix was included. Each line shows the change in mean selection accuracy over 10 cycles of crossing, testing, and selection for 100 simulation replicates. To aid visualization, the first panel shows the selection accuracies for the conventional early-stage testing strategies, and the second panel shows the selection accuracies for the sparse testing strategies and the baseline scenario.

While the selection accuracies of the sparse testing strategies were comparable to those of the conventional early-stage testing strategies, it is important to note that the sparse testing strategies evaluated a significantly higher number of genotypes using the same number of total plots, with only 10% of all genotypes replicated once (p-rep = 0.1) across the entire set of sampled test sites. In contrast, the conventional testing strategies replicated each genotype multiple times within and/or across sites. This highlights the potential of the GRM in replacing individual replication through partial replication, achieved by testing full-sibs, half-sibs, and other less closely related individuals in the breeding population. In particular, the ST-50_44 strategy demonstrated on average the highest selection accuracy, while the ST-5_440 strategy showed the lowest (Fig. S3). All three sparse testing strategies outperformed the baseline scenario in terms of accuracy across all cycles.

Additionally, compared to the conventional testing strategies with GRM, the selection accuracy in the sparse testing scenarios exhibited on average a lower standard deviation, indicating slightly greater stability in selection accuracy over cycles (Fig. S4). Notably, ST-50_44 and ST-440_5 showed a substantially lower standard deviation in accuracy than the ST-5_440 scenario. Conversely, the baseline scenario showed the highest standard deviation among all simulated scenarios.

Figure 5 a) also shows that including a GRM in the conventional early-stage testing strategies significantly increased selection accuracy. Regardless of whether a GRM was used, CT-220_5 consistently achieved the highest selection accuracy among the three conventional strategies, while CT-550_2 showed the lowest. Thus, in contrast to genetic gain, the inclusion of a GRM did not affect the ranking of the conventional strategies for selection accuracy. However, CT-220_5 exhibited the lowest increase in selection accuracy following the inclusion of the GRM, since its higher replication within and across sites, compared to the other two strategies, had already led to substantial accuracy.

Specifically, CT-440_5 and CT-550_2 showed average increases in selection accuracy of approximately 50%, while CT-220_5 showed an increase of 24%, highlighting that greater numbers of individuals can substantially improve effective replication when using the GRM. Additionally, the three conventional early-stage testing strategies with GRM exhibited higher standard deviations of selection accuracy compared to those without GRM (Fig. S4). However, due to the higher average selection accuracy when the GRM was included, coefficients of variation of selection accuracy were similar (Fig. S4). In all conventional and sparse testing simulation scenarios, selection accuracy decreased over time.

## Discussion

Early-stage sparse testing strategies can significantly increase genetic gain in plant breeding programmes compared to conventional fully replicated testing strategies. This is achieved through an efficient combination of

1. high selection accuracy, enabled by broader sampling of testing locations across the target population of environments (TPE) using a genomic relationship matrix (GRM), and
2. an increased number of tested selection candidates at a significantly reduced partial replication rate, resulting in a larger training population and a substantially higher selection intensity.

Additionally, sparse testing can lead to more stable rates of genetic gain, resulting from a more representative coverage of the TPE at early stages of testing.

While incorporating a GRM in conventional early-stage testing strategies can also enhance genetic gain, the full potential of the GRM is only realized when applied within the framework of sparse testing. This creates an opportunity to rethink early-stage testing designs in plant breeding programmes, shifting the focus from trial sites managed by breeders directly to the TPE, such as farmers’ fields, to increase the correlation between the selection environment and the TPE (Werner et al., 2025).

### Inclusion of a genomic relationship matrix in conventional early-stage testing strategies significantly enhances genetic gain

In all three conventional early-stage testing strategies, the inclusion of a GRM led to a substantial increase in genetic gain. Since all other simulation parameters remained unchanged, this improvement was exclusively driven by enhanced selection accuracy.

Genomic selection has introduced a fundamental paradigm shift in plant breeding. As the GRM uncovers genetic similarity or relatedness among individuals based on shared genomic information, such as the allelic states of SNP markers obtained from DNA microarrays, the focus is redirected from replicating the selection candidate itself to replicating the marker haplotypes (Lorenz, 2013; Lorenz and Nice, 2017) shared among full-sibs, half-sibs, and other relatives, which effectively serve as ’partial replicates’. As a result, the selection accuracy of an individual is enhanced by leveraging information from numerous other genotyped and phenotyped relatives in the population (Windhausen et al., 2012; Lorenz, 2013; Endelman et al., 2014).

Interestingly, without the use of a GRM, CT-220_5 achieved the highest genetic gain, while CT-550_2 achieved the lowest. CT-220_5 tested 220 genotypes with two replicates across five locations, whereas CT-550_2 tested two replicates in only two locations to accommodate the larger pool of 550 genotypes. Despite CT-550_2 more than doubling the number of individuals, CT-220_5 generated 15% greater genetic gain after 10 cycles, driven by over 40% higher selection accuracy. This highlights the trade-offs among accuracy, selection intensity, and genetic gain, emphasizing the need to optimize testing designs for efficient resource allocation in plant breeding programmes (Lorenz, 2013; Endelman et al., 2014; Atlin and Econopouly, 2022), while demonstrating the risk of evaluating genotypes in non-representative environments when the number of test sites is low.

Among the three conventional testing strategies with GRM, CT-440_5-GRM achieved the greatest genetic gain, facilitated by a favourable trade-off between selection intensity and accuracy. In comparison to the CT-220_5-GRM strategy, CT-440_5-GRM tested twice as many selection candidates, also across five locations, but without replication within locations. When no GRM was used, the reduced replication rate led to a decline in prediction accuracy of across-TPE performance, which had a greater adverse effect on genetic gain than the increase in selection intensity could compensate for. However, the inclusion of the GRM offset the reduction in selection accuracy, achieving nearly the same level of accuracy as CT-220_5-GRM, while still benefiting from testing more selection candidates.

Interestingly, BSLN-1_2200 achieved substantially higher genetic gain than all three conventional testing strategies, both with and without GRM. The baseline scenario represents an early generation nursery testing 2,000 genotypes at a 0.1 partial replication rate, grown at a single location. This suggests that the increase in selection intensity from evaluating four to ten times more candidates could, on average, more than offset the limited sampling of the TPE. However, the standard deviation of genetic gain across the 100 simulation replicates was significantly higher for BSLN-1_2200 than for any other testing strategy, indicating that single site selection experiments pose a substantial risk for real world breeding programmes. As further detailed in the following sections, despite its potential to deliver increased genetic gain, BSLN-1_2200 introduced considerable volatility and therefore cannot be recommended.

Including a GRM in conventional testing strategies involves relatively minimal effort and cost, yet it can have a substantial impact on genetic gain. By carefully exploring the trade-offs between selection intensity and accuracy (Atlin and Econopouly, 2022), breeders can maximize genetic gain while offsetting additional costs associated with genotyping. However, the observed increases in accuracy across the three conventional early-stage testing strategies were solely due to enhanced accuracy in the tested locations. Although including the GRM optimized the use of trial data by connecting and sharing information among selection candidates within and across tested locations, other opportunities to increase genetic gain – such as increasing selection intensity through reduced replication or expanding the number of trial sites for better representation of the TPE – remained unexploited in the conventional testing strategies.

### Early-stage sparse testing maximizes genetic gain by fully leveraging the genomic relationship matrix

The sparse testing strategies generated significantly higher genetic gain than the conventional early-stage testing strategies representing current practice in many plant breeding programmes. Increased gains reached over 20% relative to the conventional testing strategies with a GRM, and nearly 40% when no GRM was included. This improvement resulted from more efficient experimental designs and resource utilization, achieved through the optimal exploitation of the GRM, while the conventional strategies failed to leverage genomic relationship information as an opportunity to rethink early-stage testing.

Conventional testing strategies aim to achieve high selection accuracies by intensively replicating selection candidates within and across locations. While this approach is effective in the absence of genomic relationship information, as replication is the only means to improve accuracy and ensure comparability, it comes at the cost of reduced selection intensity. Although this trade-off can be optimized, the need for replication limits the potential to increase selection intensity. Incorporating a GRM, however, provides new opportunities to balance selection intensity and accuracy more effectively, as replication of marker haplotypes allows for a reduction in replication of individual selection candidates (Lorenz, 2013; Endelman et al., 2014; Montesinos-López et al., 2023). With the opportunity to increase selection intensity underutilized, conventional early-stage testing designs fall short of fully capitalizing on the GRM.

By contrast, sparse testing strategies efficiently balance the trade-off between selection intensity and accuracy to maximize genetic gain by:

1. Maximizing selection intensity through testing a substantially increased number of selection candidates at a significantly reduced partial replication rate (e.g., Lorenz, 2013).
2. Increasing selection accuracy by including a GRM to borrow information from a much larger number of relatives in the population, resulting from the increased number of selection candidates (e.g., Windhausen et al., 2012; Lorenz and Nice, 2017).
3. Enhancing selection accuracy by testing in more locations within the TPE to ensure a more representative sample of test sites (e.g., Endelman et al., 2014; Werner et al., 2024).

Thereby, sparse testing designs enabled selection accuracies comparable to those of the fully replicated conventional strategies, while testing four to ten times more selection candidates without increasing the total number of test plots.

The advantage of increasing selection intensity is particularly evident when comparing CT-220_5-GRM and CT-440_5-GRM to ST-5_440. Although all three strategies utilized a GRM and tested in five locations, ST-5_440 generated significantly greater genetic gain than the two conventional strategies, despite a slightly lower selection accuracy. Increasing the number of sparse testing locations beyond five resulted in further, albeit modest, improvements in selection accuracy, which in turn enhanced genetic gain. However, the benefits of adding more test locations are likely to be greater in years when GxE variance within the TPE is particularly high.

Conversely, reducing the number of test locations had a detrimental effect on both selection accuracy and genetic gain, as demonstrated by the baseline scenario (BSLN-1_2200). While BSLN-1_2200, on average, outperformed the conventional scenarios, both with and without a GRM due to its substantially higher selection intensity, it generated significantly less genetic gain than the sparse testing strategies. Furthermore, testing in only one location introduced a substantial risk of low or negative selection accuracies, due to occasional sampling of a test location that was not representative of the TPE. As shown in Figure 3, BSLN-1_2200 exhibited several simulation replicates with negative genetic gain in the early cycles, and overall low genetic gain in the final cycle, highlighting its inadequacy as a testing strategy for real-world breeding programmes. Given that this risk is shared by all testing strategies limited to one or very few locations, the implementation of multi-location sparse testing designs is encouraged not only for their ability to increase genetic gain on average but also for their capacity to ensure stability in genetic gain over time.

### Early-stage sparse testing strategies enhance stability of genetic gain

Sparse testing across multiple locations not only increased total genetic gain but also led to more stable rates of genetic gain over time. As the number of trial sites increased, a more representative coverage of the TPE was achieved, reducing the effect of individual sites on selection accuracy and, thereby, mitigating the impact of off-TPE locations. Hence, by incorporating more trial sites into early-stage testing, breeders can achieve both high and sustainable rates of genetic gain. To fully appreciate the benefits of early-stage sparse testing strategies, it is important to consider both the average genetic gain and the variance in genetic gain across simulation replicates.

Recall that at the start of each cycle, a new set of trial sites was randomly selected from the 2,000 simulated locations. Since off-TPE locations made up only 5-10% of the entire set of locations, most of the sampled trial sites were typically representative of TPE.

However, with fewer trial sites, the risk of selecting non-representative sites increased, occasionally leading to poor or negative prediction accuracies. Conversely, smaller samples also have an increased chance of containing only highly representative locations. As a result, selection accuracies could strongly vary across cycles.

Each testing strategy was evaluated based on 100 simulation replicates, similar to running 100 breeding programmes in parallel. As average genetic gain was calculated as the mean across all 100 replicates, fluctuations in selection accuracy across cycles were mostly averaged out, resulting in relatively stable and comparable rates of average genetic gain across all sparse testing strategies. However, ST-5_440 exhibited a considerably higher standard deviation and coefficient of variation in genetic gain compared to ST- 50_44 and ST-440_5 (Fig. 4), reflecting the increased variability across simulation runs, as shown in Figure 3. Therefore, testing across more locations helps breeders reduce the risk of making selections based on trial data that may not accurately represent the TPE. Note that this risk will increase when the proportion of off-TPE locations is high or when the genetic correlations among representative sites are lower than assumed in our simulations. Under such conditions, sparse testing becomes even more valuable, as it reduces the likelihood of over-sampling unrepresentative sites during early testing stages. This benefit is even greater when genomic selection training sets include multi- generational, multi-environment trial data (Lorenz, 2013), as supported by previous simulation studies (e.g., Gaynor et al., 2017; Werner et al., 2023).

### Finding the “optimal” early-stage sparse testing design: key takeaways and practical considerations

Breeding programmes typically aim to maximize economic returns rather than focusing solely on genetic gain. Thus, the design of early-stage trials is primarily influenced by the resource and logistical constraints of breeding programmes, as well as the capacity of their testing networks with partners and farmers to accommodate trials. Optimal testing strategies can help breeders maximize genetic gains while maintaining stable resource inputs or even reducing costs. However, what constitutes an optimal strategy will vary from one breeding programme to another.

Sparse testing strategies provide breeders with the flexibility required to design early-stage experiments that effectively balance resource efficiency and practical feasibility, while ensuring high and stable genetic gains, thus laying the groundwork for robust economic returns. Furthermore, early-stage sparse testing creates new opportunities for shifting evaluations from experimental sites managed by breeders into farmers’ fields, facilitating the assessment of selection candidates under real-world growing conditions across a wide range of locations. By incorporating on-farm sparse testing alongside on-station trials, breeders can gain valuable insights into traits such as yield, which are frequently shaped by significant GxE interactions (Atlin et al., 2001; Gaffney et al., 2015; Cooper et al., 2023; Werner et al., 2025). Particularly when on-farm management differs substantially from the environmental conditions on research stations, farm-as-incomplete-block (FAIB) approaches to sparse testing are likely to offer a robust way to account for these differences in the evaluation of selection candidates (Werner et al., 2025).

In our simulations, all sparse testing designs showed high and consistent rates of genetic gain. Notably, a relatively small number of trial sites was sufficient to effectively capture the TPE, with little improvement observed beyond 50 sites. In fact, most of the benefit from sparse testing was realised with just five sites. This finding is particularly relevant, as many breeding programmes may lack the resources or capacity to establish extensive testing networks and conduct early-stage testing on dozens or hundreds of trial sites. Nevertheless, it should be noted that while most CGIAR-led breeding networks currently conduct Stage 1 testing at only one or two sites, they usually carry out late-stage testing with national partners at more than five locations. In some large programmes, this number may exceed 30. Since establishing a new trial site is significantly more expensive than reallocating plots at an existing one, sparse testing can often be implemented cost-effectively by distributing selection candidates across sites already used for late-stage trials. In most cases, breeders are well-advised to adopt sparse testing by expanding or repurposing existing locations.

Additionally, while we used a fixed number of genotypes per site in our simulations, farm-as-incomplete-block designs may not require strict adherence to this. Allowing flexibility in the number of plots would enable the inclusion of sites of varying testing capacity, as long as the proportion of sampled farms remains representative of the TPE. Optimal, programme-specific sparse testing designs can be explored using our simulations scripts available on GitHub (https://github.com/crWerner/sparseTesting) and the R package “bsd4” (https://github.com/covaruber/bsd4).

## Statements and Declarations

### Author contributions

CRW and GNA conceived and designed the study and developed the methodology. CRW wrote and conducted the simulations and analysed the results with contributions from GCP. CRW, DGD and HG conceptualised the architecture of the TPE simulation. The first draft of the manuscript was written by CRW, and all authors commented on previous versions of the manuscript. All authors read and approved the final manuscript.

## Acknowledgments

We gratefully acknowledge Daniel Tolhurst for his invaluable help with designing and simulating the TPE.

## Code availability

The code used in the simulation study is available at https://github.com/crWerner/sparseTesting.

## Supplementary Material

**Figure S1.**
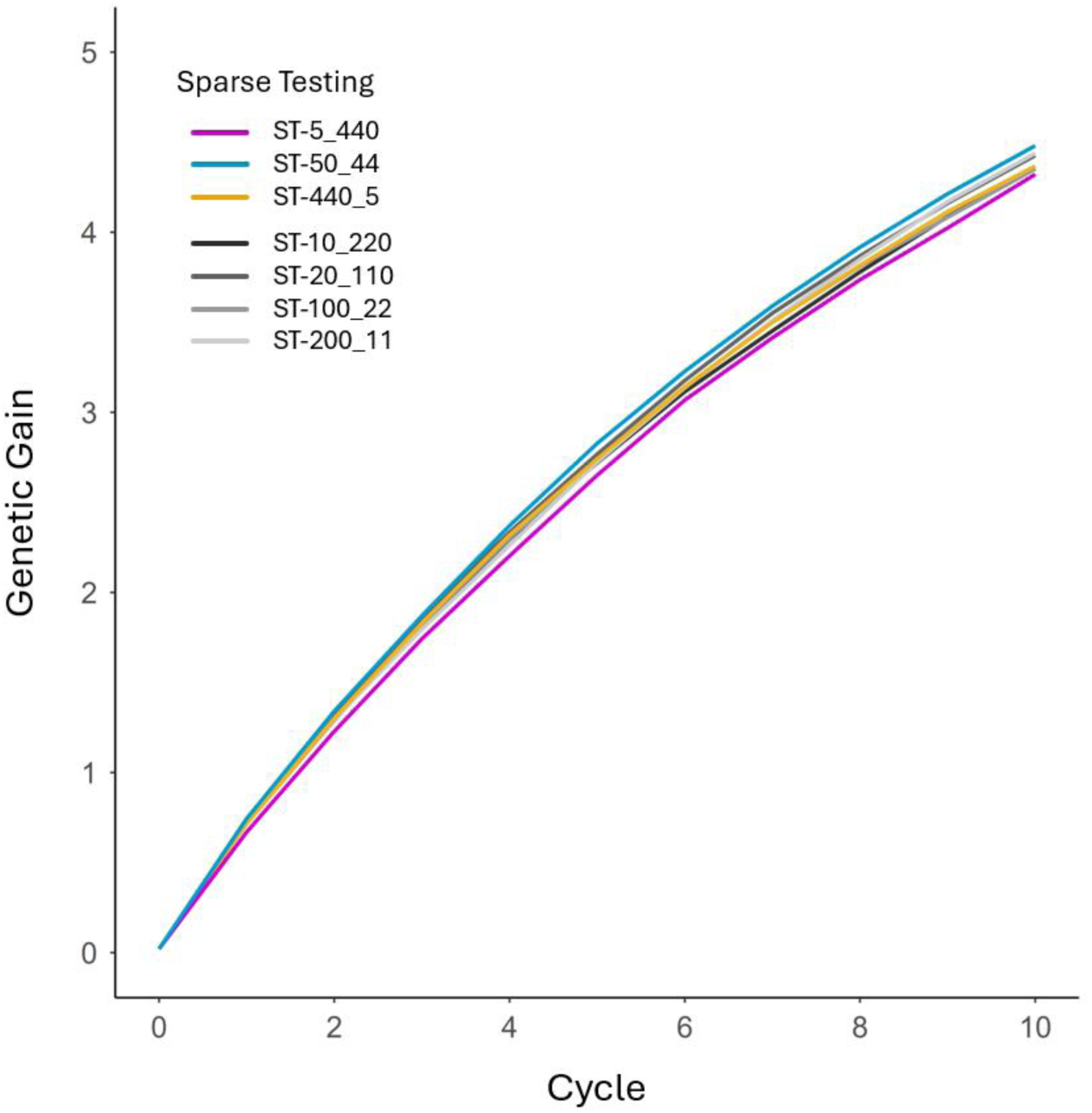
Genetic gain of the simulated conventional early-stage testing strategies without (CT-220_5, CT-440_5, CT-550_2) and with a genomic relationship matrix (CT-220_5-GRM, CT-440_5-GRM, CT-550_2-GRM), the sparse testing strategies (ST-5_440, ST-50_44, ST-440_5), and the baseline scenario (BSLN-1_2200) which represent a single-site early generation nursery including a GRM. Genetic gain is plotted as the mean genetic value of the population across all locations representative of the target population of environments. Each line shows the change in mean genetic value over 10 cycles of crossing, testing, and selection for 100 simulated replicates.

**Figure S2.**
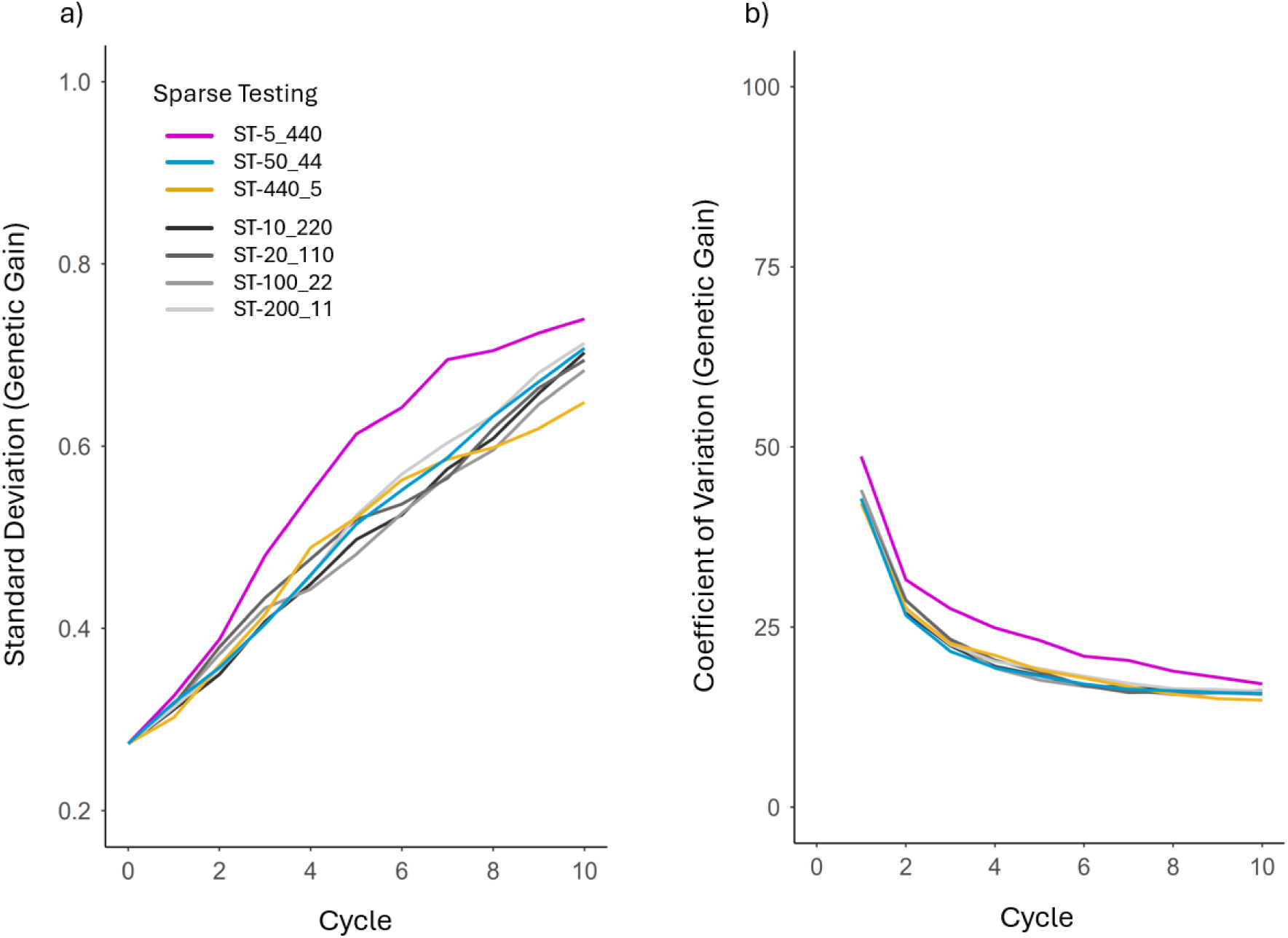
Standard deviation and coefficient of variation of genetic gain for the simulated conventional early-stage testing strategies without (CT-220_5, CT-440_5, CT-550_2) and with a genomic relationship matrix (CT-220_5-GRM, CT-440_5-GRM, CT-550_2-GRM), the sparse testing strategies (ST-5_440, ST-50_44, ST-440_5), and the baseline scenario (BSLN-1_2200). Each line shows the change in standard deviation (a) and coefficient of variation (b) over 10 cycles of crossing, testing, and selection. To aid visualization, coefficient of variation values at cycle 0 were omitted, as they ranged between 1,300 and 1,800. The standard deviation and coefficient of variation were calculated from genetic gain across the 100 simulation replicates.

**Figure S3.**
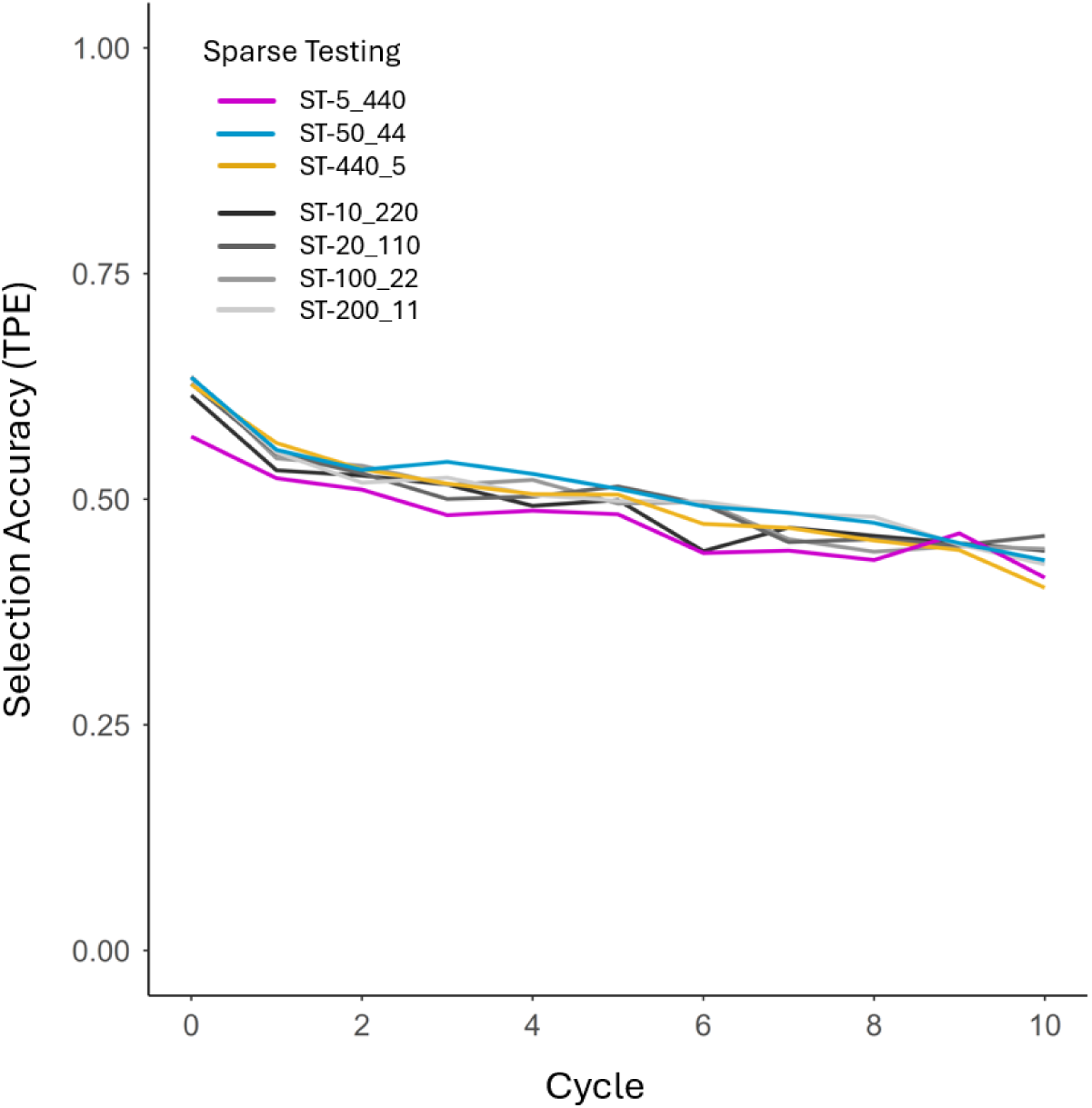
Selection accuracy for the simulated conventional early-stage testing strategies without (CT-220_5, CT-440_5, CT-550_2) and with a genomic relationship matrix (CT-220_5-GRM, CT-440_5-GRM, CT-550_2-GRM), the sparse testing strategies (ST-5_440, ST-50_44, ST-440_5), and the baseline scenario (BSLN-1_2200). Selection accuracy was calculated as the Pearson correlation coefficient between the genetic values of the genotypes across all locations representative of the target population of environments and their breeding values or genomic estimated breeding values (GEBV), depending on whether the genomic relationship matrix was included. Each line shows the change in mean selection accuracy over 10 cycles of crossing, testing, and selection for 100 simulation replicates. To aid visualization, the first panel shows the selection accuracies for the conventional early-stage testing strategies, and the second panel shows the selection accuracies for the sparse testing strategies and the baseline scenario.

**Figure S4.**
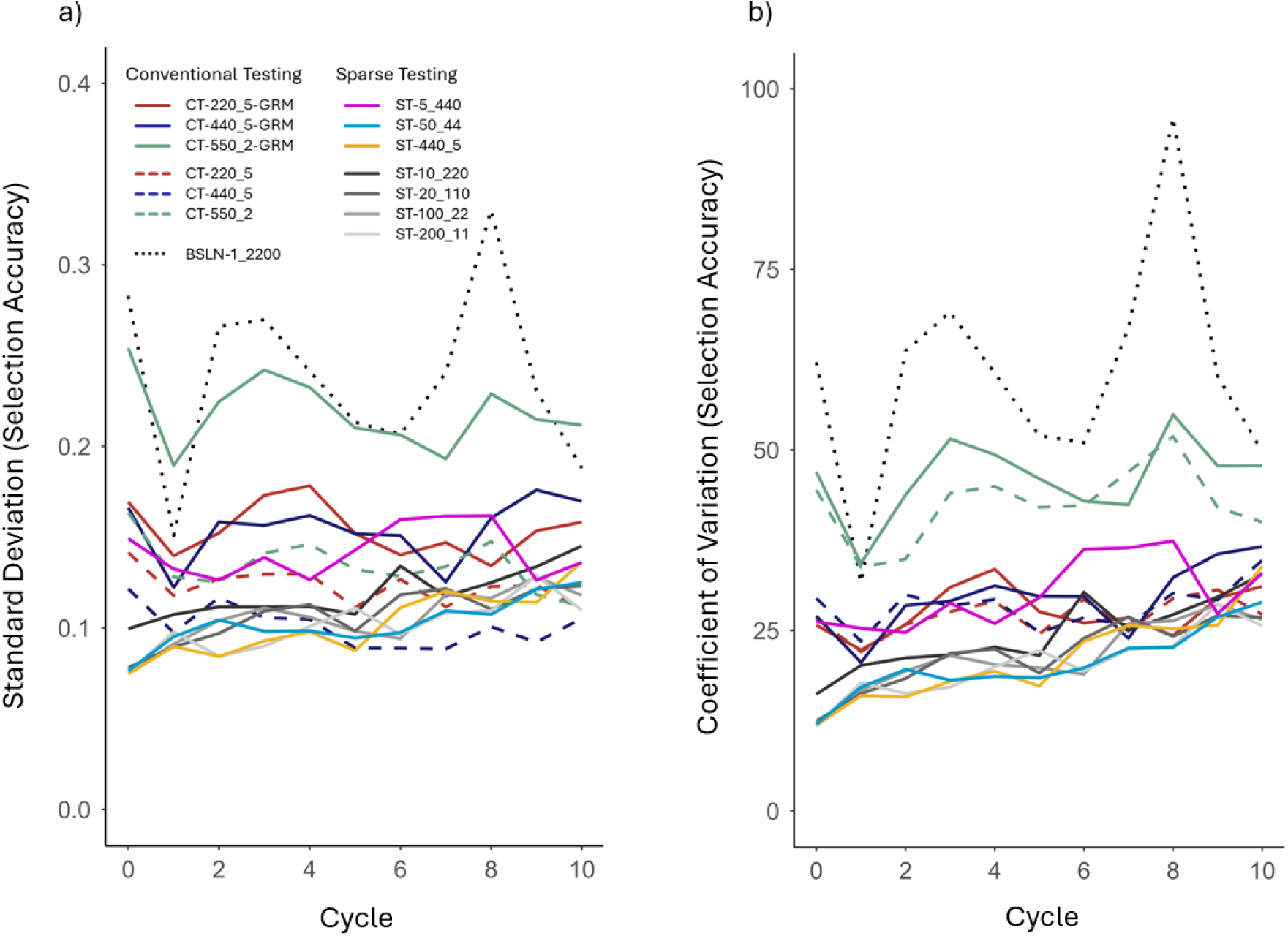
Standard deviation and coefficient of variation of genetic gain for the simulated conventional early-stage testing strategies without (CT-220_5, CT-440_5, CT-550_2) and with a genomic relationship matrix (CT-220_5-GRM, CT-440_5-GRM, CT- 550_2-GRM), the sparse testing strategies (ST-5_440, ST-50_44, ST-440_5), and the baseline scenario (BSLN-1_2200). Each line shows the change in standard deviation (a) and coefficient of variation (b) over 10 cycles of crossing, testing, and selection. To aid visualization, coefficient of variation values at cycle 0 were omitted, as they ranged between 1,300 and 1,800. The standard deviation and coefficient of variation were calculated from genetic gain across the 100 simulation replicates.

**Figure S5.**
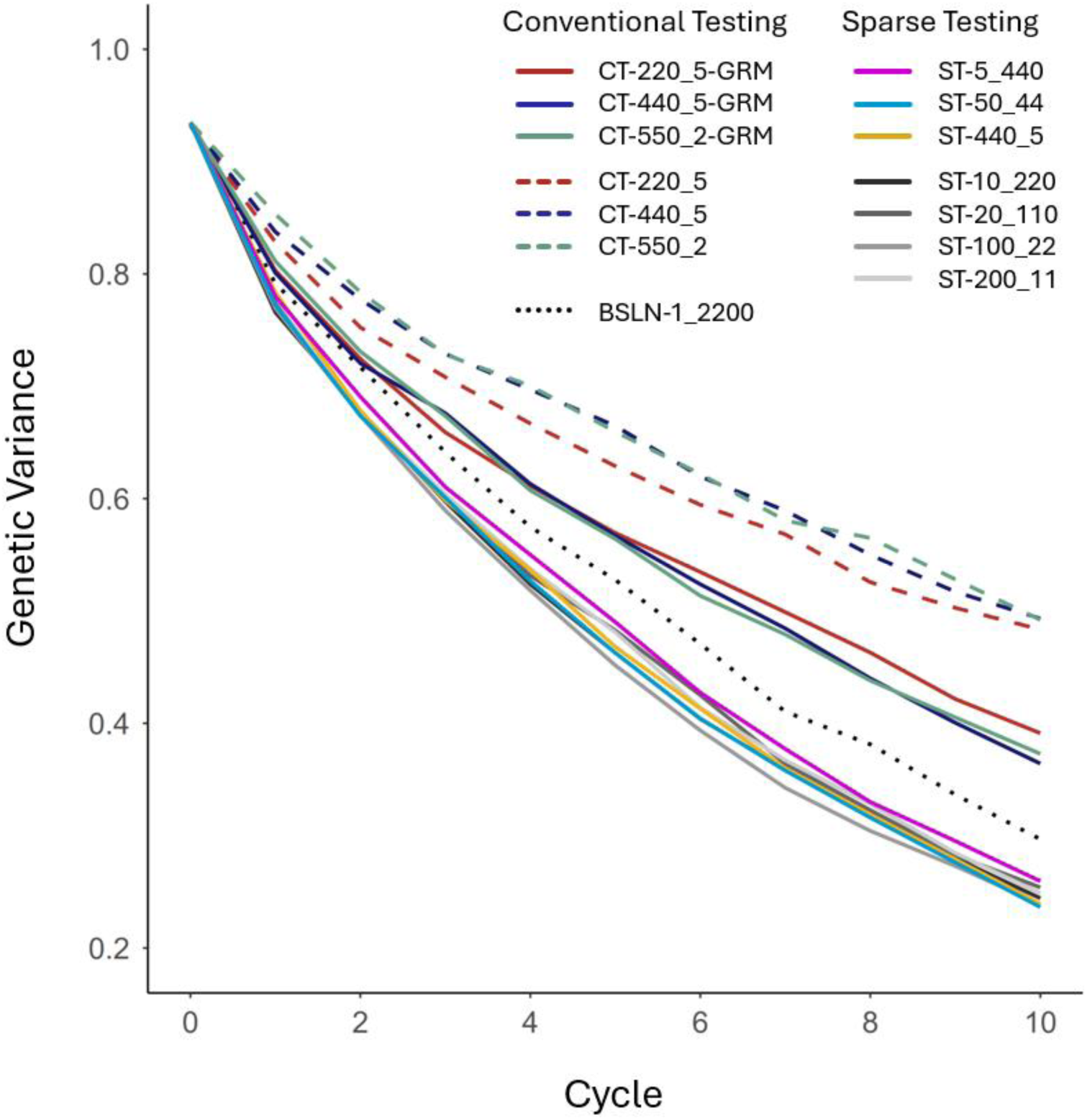
Genetic variance of the simulated conventional early-stage testing strategies without (CT-220_5, CT-440_5, CT-550_2) and with a genomic relationship matrix (CT-220_5-GRM, CT-440_5-GRM, CT-550_2-GRM), the sparse testing strategies (ST-5_440, ST-50_44, ST-440_5), and the baseline scenario (BSLN-1_2200). Genetic variance is plotted as the mean genetic variance of the population across all locations representative of the target population of environments. Each line shows the change in mean genetic variance over 10 cycles of crossing, testing, and selection for 100 simulated replicates.

**Table S1.** Summary statistics for the simulated conventional early-stage testing scenarios, all sparse testing scenarios, and the baseline scenario. The table includes average genetic gain (Gain), minimum (Min) and maximum (Max) genetic gain, standard deviation of genetic gain (SD), and coefficient of variation of genetic gain (CV), based on 100 simulation replicates.

| Scenario | Gain | Min | Max | SD | CV |
| --- | --- | --- | --- | --- | --- |
| CT-220_5 | 3.28 | 2.04 | 5.63 | 0.69 | 20.97 |
| CT-440_5 | 3.06 | 1.79 | 5.71 | 0.74 | 24.05 |
| CT-550_2 | 2.84 | 1.55 | 4.40 | 0.64 | 22.60 |
| CT-220_5-GRM | 3.51 | 1.92 | 5.21 | 0.64 | 18.21 |
| CT-440_5-GRM | 3.67 | 2.61 | 6.08 | 0.66 | 18.02 |
| CT-550_2-GRM | 3.51 | 1.31 | 5.28 | 0.73 | 20.72 |
| ST-5_440 | 4.32 | 2.71 | 6.80 | 0.74 | 17.11 |
| ST-10_220 | 4.35 | 3.02 | 6.22 | 0.70 | 16.14 |
| ST-20_110 | 4.43 | 3.06 | 6.35 | 0.69 | 15.69 |
| ST-50_44 | 4.48 | 3.20 | 6.42 | 0.71 | 15.79 |
| ST-100_22 | 4.35 | 2.85 | 6.16 | 0.68 | 15.70 |
| ST-200_11 | 4.44 | 2.94 | 6.77 | 0.71 | 16.07 |
| ST-440_5 | 4.37 | 3.03 | 6.05 | 0.65 | 14.84 |
| BSLN-1_2200 | 3.83 | 1.11 | 6.43 | 0.91 | 23.88 |

**Table S2.**
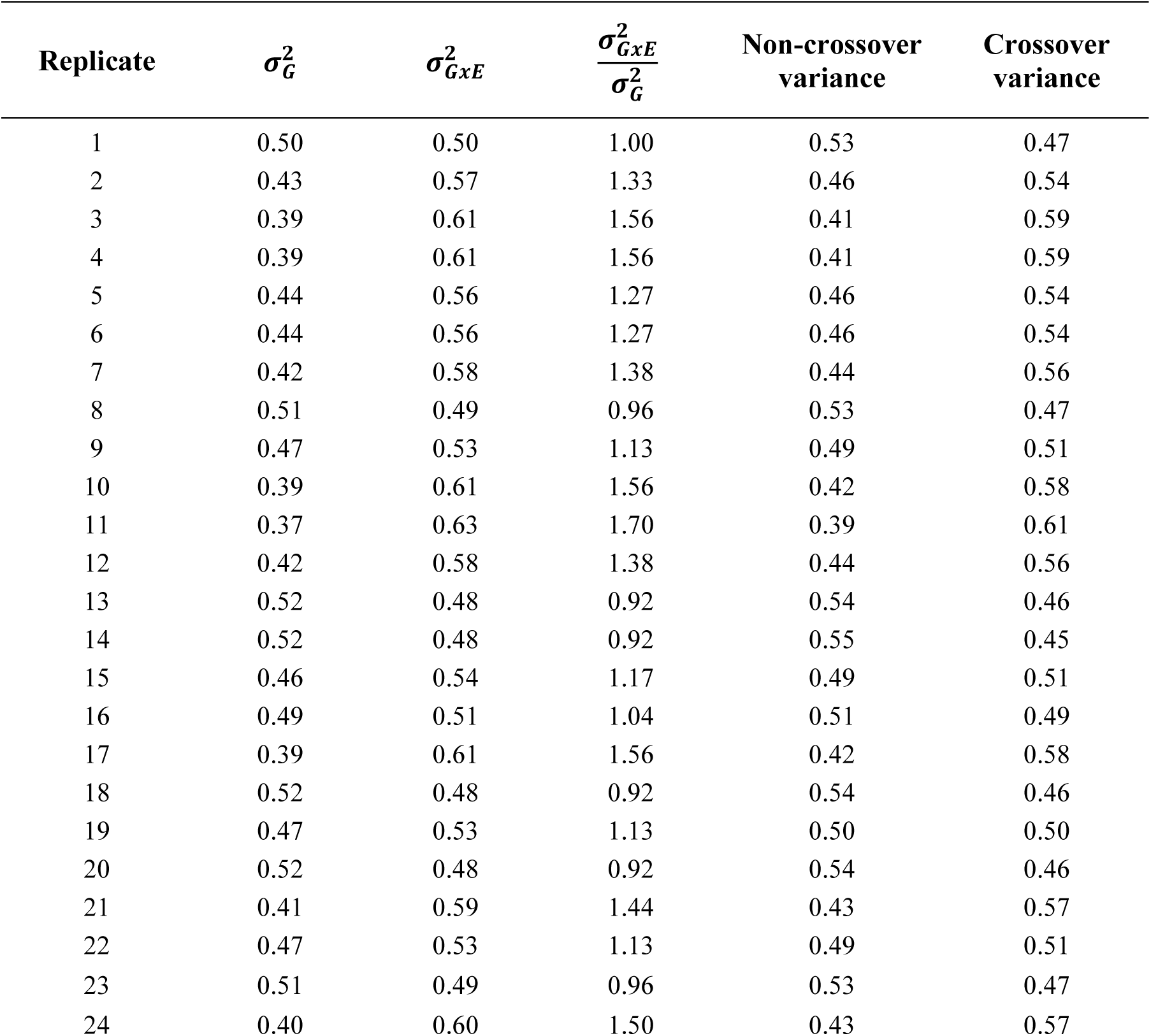
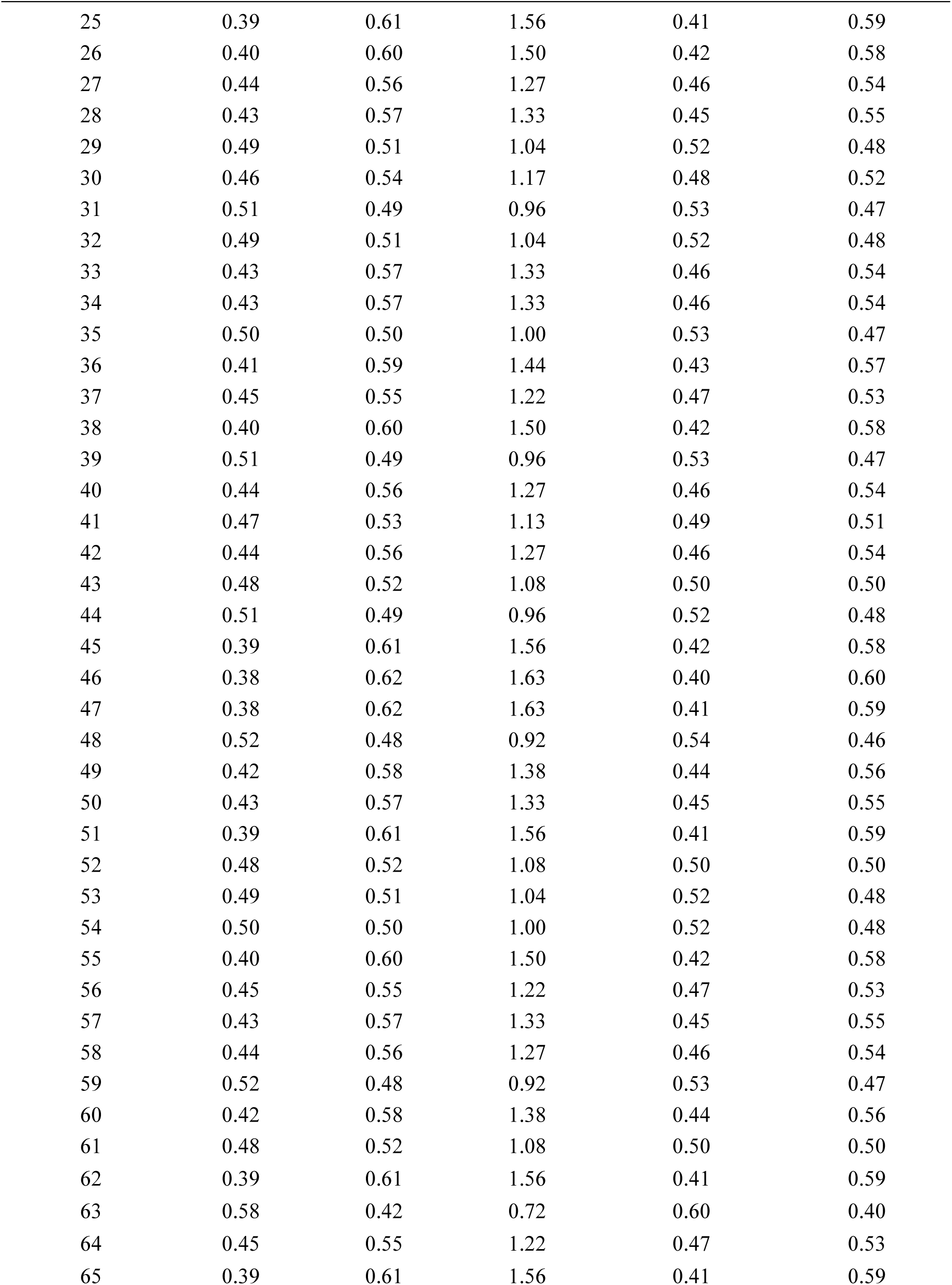
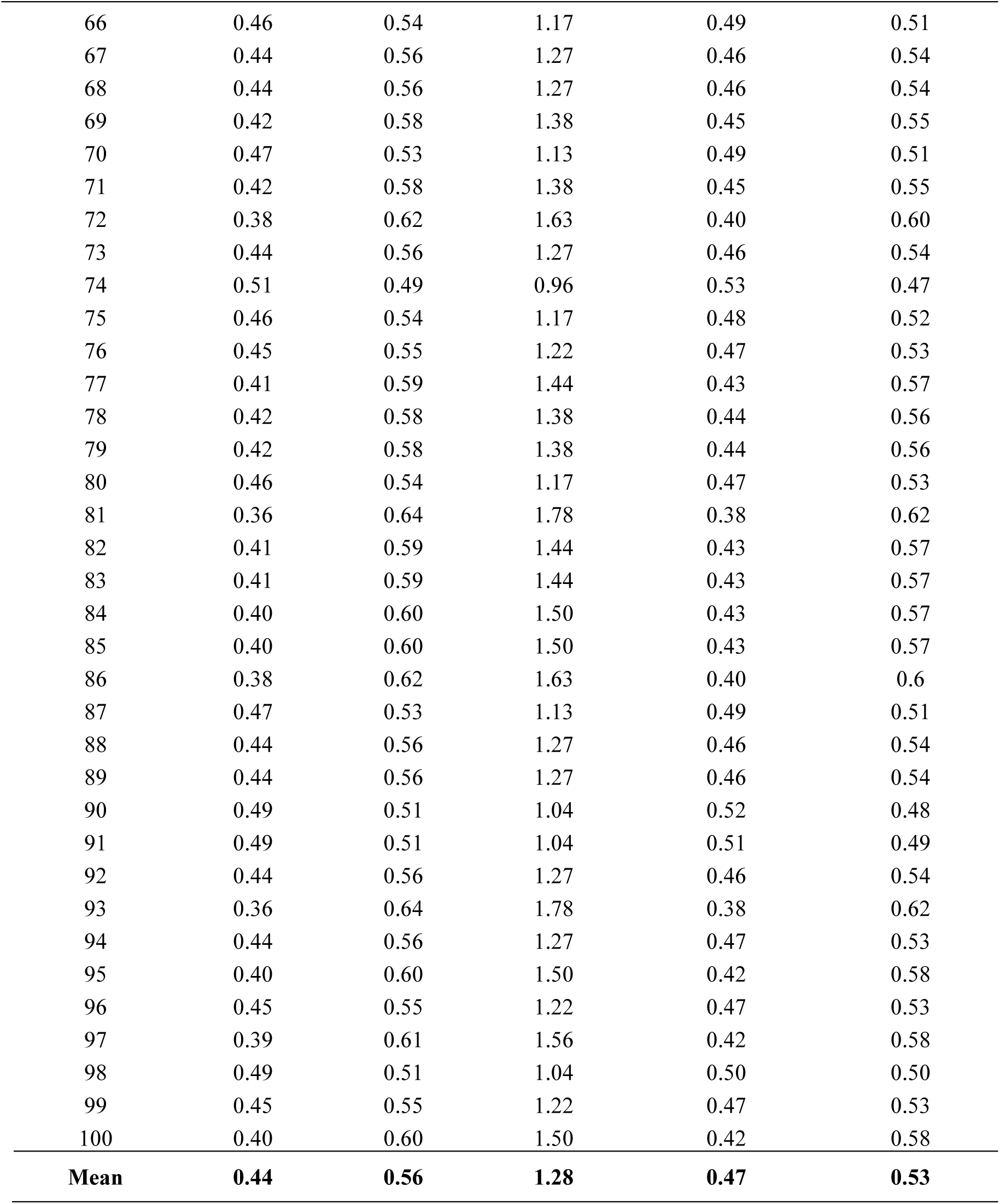
Relative proportions of main effect variance 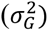 and interaction variance 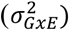 in the total genetic variance for each of the 100 simulation replicates. Variance components were derived from the genetic variance–covariance matrices in the first simulated F_2_ generation. Due to scaling of the total genetic variance to 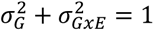, the resulting proportions are calculated as 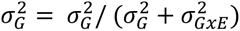 and 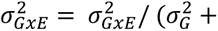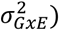., respectively. Additionally, the total genetic variance was decomposed into non-crossover and crossover variance components. All calculations were performed using the ‘*measure_variances*’ function from the R package FieldSimR (Werner et al., 2024).

| Replicate | $\sigma_G^2$ | $\sigma_{GxE}^2$ | $\frac{\sigma_{GxE}^2}{\sigma_G^2}$ | Non-crossover variance | Crossover variance |
| --- | --- | --- | --- | --- | --- |
| 1 | 0.50 | 0.50 | 1.00 | 0.53 | 0.47 |
| 2 | 0.43 | 0.57 | 1.33 | 0.46 | 0.54 |
| 3 | 0.39 | 0.61 | 1.56 | 0.41 | 0.59 |
| 4 | 0.39 | 0.61 | 1.56 | 0.41 | 0.59 |
| 5 | 0.44 | 0.56 | 1.27 | 0.46 | 0.54 |
| 6 | 0.44 | 0.56 | 1.27 | 0.46 | 0.54 |
| 7 | 0.42 | 0.58 | 1.38 | 0.44 | 0.56 |
| 8 | 0.51 | 0.49 | 0.96 | 0.53 | 0.47 |
| 9 | 0.47 | 0.53 | 1.13 | 0.49 | 0.51 |
| 10 | 0.39 | 0.61 | 1.56 | 0.42 | 0.58 |
| 11 | 0.37 | 0.63 | 1.70 | 0.39 | 0.61 |
| 12 | 0.42 | 0.58 | 1.38 | 0.44 | 0.56 |
| 13 | 0.52 | 0.48 | 0.92 | 0.54 | 0.46 |
| 14 | 0.52 | 0.48 | 0.92 | 0.55 | 0.45 |
| 15 | 0.46 | 0.54 | 1.17 | 0.49 | 0.51 |
| 16 | 0.49 | 0.51 | 1.04 | 0.51 | 0.49 |
| 17 | 0.39 | 0.61 | 1.56 | 0.42 | 0.58 |
| 18 | 0.52 | 0.48 | 0.92 | 0.54 | 0.46 |
| 19 | 0.47 | 0.53 | 1.13 | 0.50 | 0.50 |
| 20 | 0.52 | 0.48 | 0.92 | 0.54 | 0.46 |
| 21 | 0.41 | 0.59 | 1.44 | 0.43 | 0.57 |
| 22 | 0.47 | 0.53 | 1.13 | 0.49 | 0.51 |
| 23 | 0.51 | 0.49 | 0.96 | 0.53 | 0.47 |
| 24 | 0.40 | 0.60 | 1.50 | 0.43 | 0.57 |

| Replicate | $\sigma_G^2$ | $\sigma_{G \times E}^2$ | $\frac{\sigma_{G \times E}^2}{\sigma_G^2}$ | Non-crossover<br>variance | Crossover<br>variance |
| --- | --- | --- | --- | --- | --- |
| 25 | 0.39 | 0.61 | 1.56 | 0.41 | 0.59 |
| 26 | 0.40 | 0.60 | 1.50 | 0.42 | 0.58 |
| 27 | 0.44 | 0.56 | 1.27 | 0.46 | 0.54 |
| 28 | 0.43 | 0.57 | 1.33 | 0.45 | 0.55 |
| 29 | 0.49 | 0.51 | 1.04 | 0.52 | 0.48 |
| 30 | 0.46 | 0.54 | 1.17 | 0.48 | 0.52 |
| 31 | 0.51 | 0.49 | 0.96 | 0.53 | 0.47 |
| 32 | 0.49 | 0.51 | 1.04 | 0.52 | 0.48 |
| 33 | 0.43 | 0.57 | 1.33 | 0.46 | 0.54 |
| 34 | 0.43 | 0.57 | 1.33 | 0.46 | 0.54 |
| 35 | 0.50 | 0.50 | 1.00 | 0.53 | 0.47 |
| 36 | 0.41 | 0.59 | 1.44 | 0.43 | 0.57 |
| 37 | 0.45 | 0.55 | 1.22 | 0.47 | 0.53 |
| 38 | 0.40 | 0.60 | 1.50 | 0.42 | 0.58 |
| 39 | 0.51 | 0.49 | 0.96 | 0.53 | 0.47 |
| 40 | 0.44 | 0.56 | 1.27 | 0.46 | 0.54 |
| 41 | 0.47 | 0.53 | 1.13 | 0.49 | 0.51 |
| 42 | 0.44 | 0.56 | 1.27 | 0.46 | 0.54 |
| 43 | 0.48 | 0.52 | 1.08 | 0.50 | 0.50 |
| 44 | 0.51 | 0.49 | 0.96 | 0.52 | 0.48 |
| 45 | 0.39 | 0.61 | 1.56 | 0.42 | 0.58 |
| 46 | 0.38 | 0.62 | 1.63 | 0.40 | 0.60 |
| 47 | 0.38 | 0.62 | 1.63 | 0.41 | 0.59 |
| 48 | 0.52 | 0.48 | 0.92 | 0.54 | 0.46 |
| 49 | 0.42 | 0.58 | 1.38 | 0.44 | 0.56 |
| 50 | 0.43 | 0.57 | 1.33 | 0.45 | 0.55 |
| 51 | 0.39 | 0.61 | 1.56 | 0.41 | 0.59 |
| 52 | 0.48 | 0.52 | 1.08 | 0.50 | 0.50 |
| 53 | 0.49 | 0.51 | 1.04 | 0.52 | 0.48 |
| 54 | 0.50 | 0.50 | 1.00 | 0.52 | 0.48 |
| 55 | 0.40 | 0.60 | 1.50 | 0.42 | 0.58 |
| 56 | 0.45 | 0.55 | 1.22 | 0.47 | 0.53 |
| 57 | 0.43 | 0.57 | 1.33 | 0.45 | 0.55 |
| 58 | 0.44 | 0.56 | 1.27 | 0.46 | 0.54 |
| 59 | 0.52 | 0.48 | 0.92 | 0.53 | 0.47 |
| 60 | 0.42 | 0.58 | 1.38 | 0.44 | 0.56 |
| 61 | 0.48 | 0.52 | 1.08 | 0.50 | 0.50 |
| 62 | 0.39 | 0.61 | 1.56 | 0.41 | 0.59 |
| 63 | 0.58 | 0.42 | 0.72 | 0.60 | 0.40 |
| 64 | 0.45 | 0.55 | 1.22 | 0.47 | 0.53 |
| 65 | 0.39 | 0.61 | 1.56 | 0.41 | 0.59 |
| 66 | 0.46 | 0.54 | 1.17 | 0.49 | 0.51 |
| 67 | 0.44 | 0.56 | 1.27 | 0.46 | 0.54 |
| 68 | 0.44 | 0.56 | 1.27 | 0.46 | 0.54 |
| 69 | 0.42 | 0.58 | 1.38 | 0.45 | 0.55 |
| 70 | 0.47 | 0.53 | 1.13 | 0.49 | 0.51 |
| 71 | 0.42 | 0.58 | 1.38 | 0.45 | 0.55 |
| 72 | 0.38 | 0.62 | 1.63 | 0.40 | 0.60 |
| 73 | 0.44 | 0.56 | 1.27 | 0.46 | 0.54 |
| 74 | 0.51 | 0.49 | 0.96 | 0.53 | 0.47 |
| 75 | 0.46 | 0.54 | 1.17 | 0.48 | 0.52 |
| 76 | 0.45 | 0.55 | 1.22 | 0.47 | 0.53 |
| 77 | 0.41 | 0.59 | 1.44 | 0.43 | 0.57 |
| 78 | 0.42 | 0.58 | 1.38 | 0.44 | 0.56 |
| 79 | 0.42 | 0.58 | 1.38 | 0.44 | 0.56 |
| 80 | 0.46 | 0.54 | 1.17 | 0.47 | 0.53 |
| 81 | 0.36 | 0.64 | 1.78 | 0.38 | 0.62 |
| 82 | 0.41 | 0.59 | 1.44 | 0.43 | 0.57 |
| 83 | 0.41 | 0.59 | 1.44 | 0.43 | 0.57 |
| 84 | 0.40 | 0.60 | 1.50 | 0.43 | 0.57 |
| 85 | 0.40 | 0.60 | 1.50 | 0.43 | 0.57 |
| 86 | 0.38 | 0.62 | 1.63 | 0.40 | 0.6 |
| 87 | 0.47 | 0.53 | 1.13 | 0.49 | 0.51 |
| 88 | 0.44 | 0.56 | 1.27 | 0.46 | 0.54 |
| 89 | 0.44 | 0.56 | 1.27 | 0.46 | 0.54 |
| 90 | 0.49 | 0.51 | 1.04 | 0.52 | 0.48 |
| 91 | 0.49 | 0.51 | 1.04 | 0.51 | 0.49 |
| 92 | 0.44 | 0.56 | 1.27 | 0.46 | 0.54 |
| 93 | 0.36 | 0.64 | 1.78 | 0.38 | 0.62 |
| 94 | 0.44 | 0.56 | 1.27 | 0.47 | 0.53 |
| 95 | 0.40 | 0.60 | 1.50 | 0.42 | 0.58 |
| 96 | 0.45 | 0.55 | 1.22 | 0.47 | 0.53 |
| 97 | 0.39 | 0.61 | 1.56 | 0.42 | 0.58 |
| 98 | 0.49 | 0.51 | 1.04 | 0.50 | 0.50 |
| 99 | 0.45 | 0.55 | 1.22 | 0.47 | 0.53 |
| 100 | 0.40 | 0.60 | 1.50 | 0.42 | 0.58 |
| **Mean** | **0.44** | **0.56** | **1.28** | **0.47** | **0.53** |

